# Chemoproteomic-enabled characterization of small GTPase Rab1a as a target of an *N*-arylbenzdiimidazole ligand’s rescue of Parkinson’s-associated cell toxicity

**DOI:** 10.1101/2021.05.05.442776

**Authors:** A. Katherine Hatstat, Baiyi Quan, Morgan Bailey, Michael C. Fitzgerald, Michaela C. Reinhart, Dewey G. McCafferty

## Abstract

The development of phenotypic models of Parkinson’s disease (PD) has enabled screening and identification of phenotypically active small molecules that restore complex biological pathways affected by PD toxicity. While these phenotypic screening platforms are powerful, they do not inherently enable direct identification of the cellular targets of promising lead compounds. To overcome this, chemoproteomic platforms like Thermal Proteome Profiling (TPP) and Stability of Proteins from Rates of Oxidation (SPROX) can be implemented to reveal protein targets of biologically active small molecules. Here we utilize both of these chemoproteomic strategies to identify targets of an *N-*arylbenzdiimidazole compound, NAB2, which was previously identified for its ability to restore viability in cellular models of PD-associated α-synuclein toxicity. The combined results from our TPP and SPROX analyses of NAB2 and the proteins in a neuroblastoma-derived SHSY5Y cell lysate reveal a previously unrecognized protein target of NAB2. This newly recognized target, Rab1a, is a small GTPase that acts as a molecular switch to regulate ER-to-Golgi trafficking, a process that is disrupted by α-synuclein toxicity and restored by NAB2 treatment. Further validation reveals that NAB2 binds to Rab1a with selectivity for its GDP-bound form and that NAB2 treatment phenocopies Rab1a overexpression in alleviation of α-synuclein toxicity. Finally, we conduct a preliminary investigation into the relationship between Rab1a and the E3 ubiquitin ligase, Nedd4, a previously identified NAB2 target. Together, these efforts expand our understanding of the mechanism of NAB2 in the alleviation of α-synuclein toxicity and reinforce the utility of chemoproteomic identification of the targets of phenotypically active small molecules that regulate complex biological pathways.

## Introduction

Parkinson’s disease (PD) is a debilitating neurodegenerative disorder for which there are no neuroprotective treatments currently available to patients. While there are pharmacological approaches like dopamine replacement therapy and surgical interventions like deep brain stimulation,^1,2^ there is an urgent need for development of new therapeutics that slow down or halt the progression of the degenerative disease. To this end, there have been extensive efforts towards developing experimental models of PD-associated toxicity that can be implemented for phenotype-driven, high-throughput screening to identify novel lead compounds.^3^ The availability of these screening platforms provides a powerful tool for drug development and has enabled the identification of several promising leads for the treatment of Parkinsonian toxicity.^4–7^ There are, however, limitations in these approaches as they do not inherently enable identification of the target or mechanism of action (MOA) of the lead compound. While knowledge of MOA is not required for approval by the Food and Drug Administration (FDA), it is beneficial information that enables use of lead compounds as probes to interrogate relevant biological processes, contributing to the process of drug development.^8^ This is particularly pertinent in the case of PD as it shares many underlying mechanisms with other neurodegenerative diseases.^9,10^ Further, some phenotypically active small molecules identified to date show activity in more than one neurodegeneration model,^5,6,11–13^ suggesting target proteins or pathways that are conserved across neurodegeneration. Therefore, identifying the target of a drug and understanding its MOA may provide access to a lead that is applicable to neurodegenerative disorders more broadly.

Recently, a variety of unbiased mass spectrometry-based proteomic strategies have been developed to identify the protein targets of biologically active ligands by quantitatively measuring ligand-induced changes in protein folding stability. These strategies have included the thermal <u>p</u>roteome <u>p</u>rofiling (TPP) technique,^14^ several proteolysis-based techniques (e.g., DARTS,^15,16^ LiP,^17,18^ and <u>p</u>ulse <u>p</u>roteolysis (PP)^19–21^), and the <u>s</u>tability of <u>p</u>roteins from rates of <u>o</u>xidation (SPROX) methodology.^22–24^ These techniques complement the use of phenotype-driven high-throughput screens by enabling characterization of target engagement without chemical derivatization of the target ligand for covalent capture or enrichment. This process provides crucial information for revealing the MOA of promising lead compounds.

As a demonstration of the power of these proteomic tools in target identification of neurodegenerative disease treatment, we report here the combined use of TPP and SPROX to identify targets of a lead compound previously found through a phenotype-driven screen of PD-associated toxicity. In this case, the lead compound, which contains a *N*-arylbenzdiimidazole (NAB) scaffold, was identified for its ability to alleviate phenotypic markers of cellular stress in yeast models of TDP-43 toxicity (associated with amyotrophic lateral sclerosis, ALS) and then in models of α-synuclein toxicity (associated with PD) (**Figure 1A**).^6^ α-Synuclein is a trafficking protein in which PD-associated mutations, duplications, or triplications at *SNCA*, the gene locus encoding α-synuclein, result in expression of a toxic form of the protein that is prone to aggregation.^7,25–28^ Accumulation of toxic α-synuclein proteoforms results in cellular dysfunctions including disruption of ER-to-Golgi trafficking, formation of α-synuclein aggregates, and generation of reactive oxygen species. Structure activity relationship studies afforded improved derivative NAB2, which significantly alleviated these markers of toxicity, and counter genetic screening revealed that its activity was primarily dependent upon an E3 ubiquitin ligase, Rsp5.^6^ Further screening revealed a small network of Rsp5-associated proteins upon which the activity of the NAB scaffold was partially dependent (**Figure 1B**).^6^ Subsequent validation in iPSC-derived neuron models revealed that NAB activity was retained and was dependent upon Nedd4, the mammalian homolog of Rsp5.^6,7^ Despite identification of Nedd4 as a putative NAB target, our recent characterization of NAB2 and Nedd4 revealed that NAB2, the highest activity derivative, binds to Nedd4 *in vitro* but does not alter its *in vitro* activity, conformation, or ubiquitin linkage specificity.^29^ These results, in combination with the initial identification of a Rsp5/Nedd4 associated network implicated in the NAB mechanism, indicate that the phenotypic rescue of α-synuclein toxicity by NAB may involve multiple protein targets.

**Figure 1:**
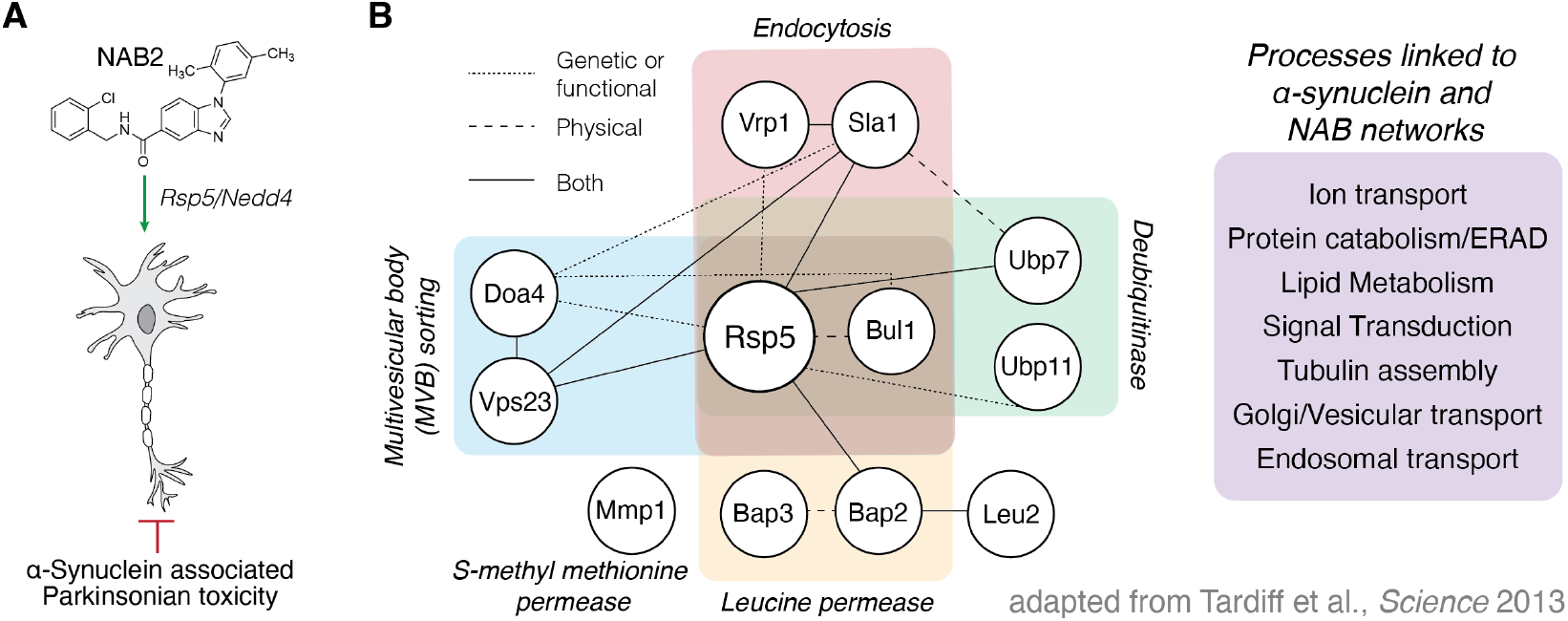
Previous efforts toward the discovery and characterization of the NAB scaffold in the rescue of α-synuclein toxicity have been Nedd4 centric but indicate other proteins are implicated in the NAB mechanism of action. **(A)** *N-*arylbenzdiimidazole compound NAB2 has been previously studied for its ability to restore phenotypic viability in experimental models of α-synuclein associated Parkinsonian toxicity. **(B)** Tardiff and co-workers^6^ in the Lindquist lab discovered the NAB scaffold in a yeast-based screen of α-synuclein toxicity and demonstrated that activity was conserved in mammalian models. Activity of the NAB scaffold was dependent upon E3 ubiquitin ligase Rsp5/Nedd4, the yeast and mammalian homologues, respectively,^6,7^ but revealed a network of proteins that were also identified in chemical genetic screening to be affected by NAB2 treatment and α-synuclein toxicity.

To investigate this, we sought an unbiased approach to identify NAB targets across the proteome to further understand the MOA of this phenotypically promising lead. To this end, we employed a chemoproteomic strategy in which a multiplexed, one-pot approach enabled parallel TPP and SPROX identification of proteins that exhibit NAB2-dependent shifts in stability.^30^ Through this analysis, we identified the small GTPase, Rab1a, as a hit in both TPP and SPROX analyses, providing Rab1a as a putative, previously unrecognized target of NAB2. The identification of Rab1a as a target in the rescue of α-synuclein toxicity is especially interesting as Rab1a regulates trafficking from the ER to Golgi, a process disrupted by α-synuclein toxicity but rescued by NAB2 in phenotypic analyses. Further, Rab1 overexpression is shown here to restore viability in models of α-synuclein toxicity, a result which is consistent with the effect of NAB2 treatment. Additional validation experiments on purified Rab1a revealed that the thermal stability of the protein is decreased by NAB2 and that binding occurs in a nucleotide-dependent manner and is selective for the GDP-bound conformation of Rab1a. Further, *in vitro* and cell-based experiments reveal that NAB2 does not alter Rab1a GTPase activity, but NAB2 treatment and Rab1a overexpression restore cell viability in the presence of α-synuclein toxicity. Finally, we explore putative links between previously studied target Nedd4 and Rab1a through cellular assays and interactome analyses. Together, these analyses provide insight into the mechanism of NAB2 as a promising phenotypically active small molecule in the rescue of trafficking defects induced by α-synuclein toxicity and provide new evidence for the role of Rab1a as a target in NAB2 activity.

## Results and discussion

### One-pot chemoproteomic method enables screening of NAB2-induced changes in protein stability via two orthogonal methods in parallel

To identify NAB2-dependent shifts in protein stability across the proteome, we employed a parallel approach using both TPP and SPROX, which measure ligand-induced changes in the thermal and chemical denaturation properties of proteins, respectively (**Figure 2**). To employ these techniques for identification of NAB2 protein targets with high confidence and in an efficient manner, a highly multiplexed “one-pot” method was used. This method, which was recently validated by Cabrera and co-workers,^30^ uses cellular lysate in the presence or absence of small molecule treatment (**Figure 2C**). The lysate samples are processed according to standard TPP or SPROX workflows, where the samples (five biological replicates of each NAB2- or vehicle-treated control) are exposed to a denaturation gradient (temperature for TPP and urea for SPROX) followed by chemical oxidation (SPROX) or ultra-centrifugation (TPP). Samples across the gradient in each replicate are pooled, and the pooled samples are submitted to a quantitative bottom-up proteomics analysis using isobaric mass tags (i.e., a TMT 10-plex labeling).^18^ The isobaric mass tag labelling strategy allows for five replicates of each experimental condition (small molecule and vehicle control) to be analyzed by LC-MS/MS using a single sample per method. Though the one-pot approach precludes generation of full denaturation curves for proteins identified, it enables high confidence identification of hit proteins through a high number of biological replicates per method, which minimizes not only the experimental cost but also the false positive rate.^30^ Requiring hit proteins to appear in multiple orthogonal methods (e.g., TPP and SPROX) also has been shown in model studies to result in false positive rates close to 0.^30^

**Figure 2:**
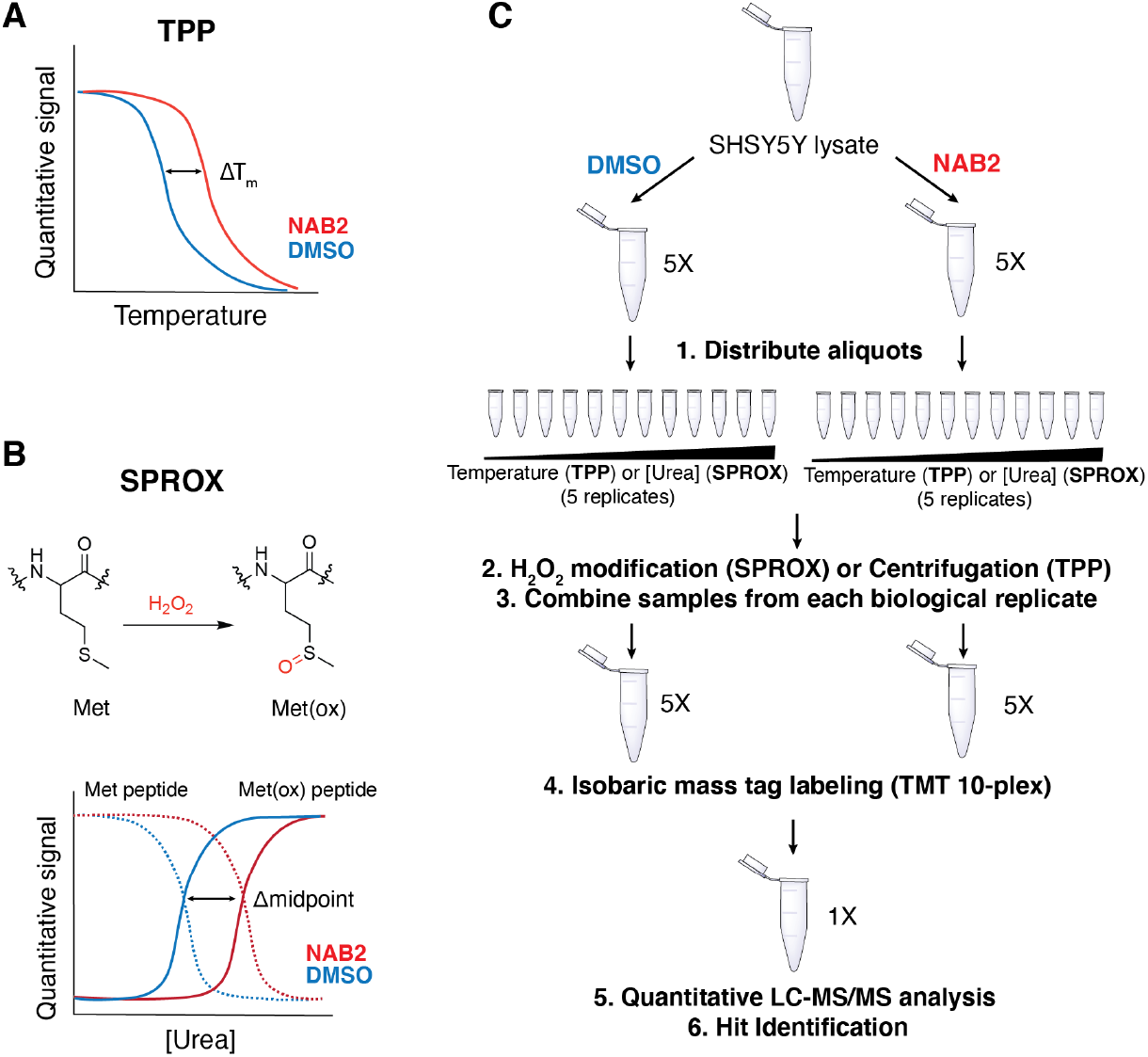
TPP and SPROX methods enable chemoproteomic analysis of ligand-dependent changes in protein stability and can be employed in parallel. **(A)** TPP measures shifts in protein thermostability as a function of melting temperature (T_m_) or the temperature at which half of the protein population is denatured. Ligand binding can thermodynamically shift protein stability and is detectable as ΔT_m_. **(B)** SPROX measures the stability of proteins as a function of the protein susceptibility to chemical oxidation over a chemical denaturation gradient. In SPROX, methionine residues are specifically oxidized in the presence of H_2_O_2_, and shifts in protein stability are measured as a function of changes in the midpoint of methionine oxidation curves. **(C)** Cabrera and co-workers recently reported a one-pot workflow for combinatorial analysis of two orthogonal chemoproteomic experiments in a single sample for LC-MS/MS analysis.^30^ This workflow was employed with TPP/SPROX to identify NAB2 targets.

### Rab1a identified as a hit in one-pot TPP/SPROX method

Using the one-pot TPP/SPROX workflow described above (**Figure 2C**), lysates of neuroblastoma-derived SHSY5Y cells were treated with NAB2 (20 µM) or DMSO and pre-incubated prior to processing for TPP/SPROX analysis. NAB2-dependent stability changes in the TPP and SPROX experiments were identified using student’s two tailed t-test and visualized by volcano plots, where the change in protein stability (measured by Z-score) in NAB2-treated samples relative to DMSO control samples is plotted against the statistical significance of the Z-score across replicates (–log(p-value)) (**Figure 3A,B**; **Supplemental Table 1**). This allows for identification of proteins that have significantly altered stabilities in response to NAB2 treatment. Hit proteins in each strategy were selected as those with a Z-score less than –2 or greater than 2 and a p-value of < 0.001 (–log(p-value) > 3.0). These thresholds were determined in the initial validation of the one-pot method as they were shown to minimize false positives.^30^ Using these thresholds, six and two proteins were identified as significant hits in the SPROX and TPP analyses (respectively), which provided for the analysis of 2174 and 2369 proteins (respectively) in the SHSY5Y cell lysates (**Figure 3**). These hit proteins were identified as: keratin (P35527), exportin-5 (Q9HAV4), bifunctional purine biosynthesis protein ATIC (P31939), coiled-coiled domain containing protein 124 (Q96CT7), annexin A6 (P08133-1) and Rab1a (P62820) for SPROX (**Figure 3A**) and proteasome adapter and scaffold protein ECM29 (Q5VYK3) and Rab1a (P62820) for TPP (**Figure 3B**). Rab1a was the only protein identified as a hit in both the TPP and SPROX experiments (**Figure 3C**). The result provides high confidence that Rab1a is a target of NAB2. The TPP result indicates the Rab1a protein is significantly destabilized (i.e., thermally denatured and precipitated at a lower temperature) in the presence of NAB2. The oxidized-methionine-containing peptide hit detected for Rab1 in the SPROX experiment indicated that the protein is chemically denatured at a higher denaturant concentration in the presence of NAB2. The chemical denaturation behavior is consistent with a direct binding interaction between NAB2 and Rab1a. The reduced melting temperature for Rab1 in the presence of NAB2 in the TPP experiment is consistent with the NAB2-induced disruption of protein-protein interactions involving Rab1 and/or with NAB2 binding inducing conformational changes in Rab1, as have been observed in PTSA studies of other protein-ligand systems.^31,32^ The results of our PTSA studies of NAB2 biding a purified construct of Rab1 suggest the latter (see below). While additional structural studies are needed to elucidate the detailed molecular basis of the NAB2-Rab1 interaction, our identification of Rab1a as a hit protein in these experiments clearly implicates this protein in NAB2’s mechanism of action.

**Figure 3:**
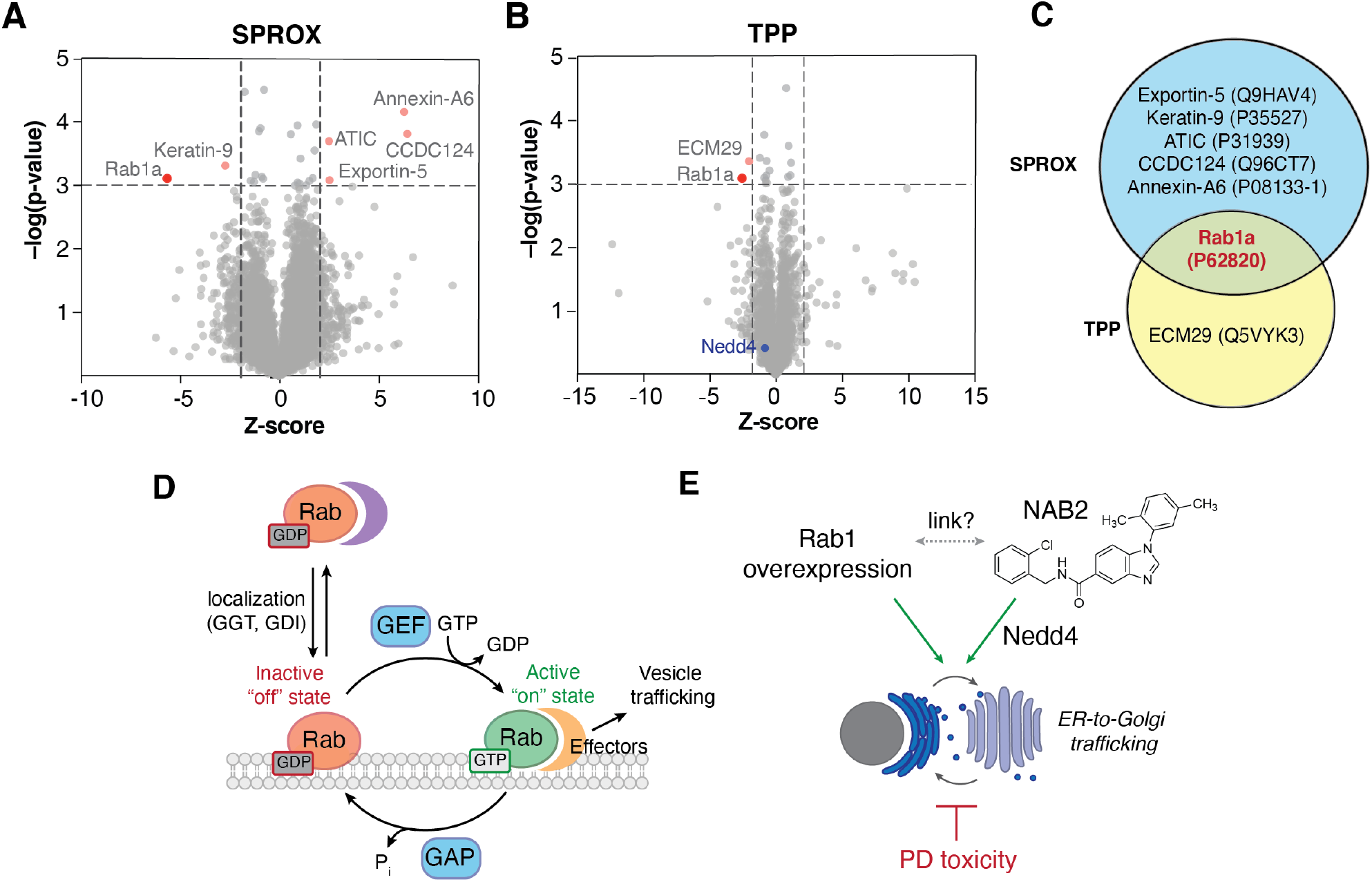
Volcano plot analysis enables identification of proteins that exhibit significant shifts in stability in response to NAB2 treatment. Protein abundances were measured quantitatively by LC-MS/MS and multiplexed data (via TMT labeling) was deconvoluted prior to volcano plot analysis of **(A)** methionine-containing peptide abundances in SPROX and **(B)** protein abundances in TPP. In both **A** and **B**, significance thresholds were defined as Z-score less than –2 or greater than 2 and a p-value of < 0.001 (–log(p-value) > 3.0). **(C)** Overalap of significantly altered proteins across TPP/SPROX experiments reveals Rab1a as a conserved NAB2-dependent hit in both experiments. **(B)** Rab GTPases regulate trafficking by acting as molecular switches that adopt distinct conformations in a nucleotide-dependent manner. Rab activity is tightly controlled by protein-protein interactions with regulatory proteins and effectors.^33,34^ GGT: geranylgeranyl transferase; GDI: guanine nucleotide dissociation factor; GEF: guanine nucleotide exchange factor; GAP: GTPase activating protein. **(C)** ER-to-Golgi trafficking is disrupted in PD-associated α-synuclein toxicity but is stimulated by NAB2 treatment^6^ and by Rab1 overexpression.^35,36^ Despite this, a link between NAB2 and Rab1 has not been previously established.

### Previously identified target Nedd4 is not identified as NAB2-dependent hit in one-pot TPP/SPROX experiment

While we have previously demonstrated that NAB2 binds to Nedd4 *in vitro*,^29^ we did not identify Nedd4 as a hit in this experiment. Nedd4-derived tryptic peptides were only identified in the TPP experiment and not in the SPROX dataset, thereby precluding it from identification as a hit in the current study. One limitation of the SPROX experiment is that it requires the detection and quantitation of methionine-containing peptides from potential protein targets in the bottom-up proteomics readout. While there are methionine residues in Nedd4 (16–20, depending on the Nedd4 isoform), methionine-containing tryptic peptides from Nedd4 were either insufficiently enriched and/or insufficiently ionized in the LC-MS/MS readout used in the SPROX experiment. Nedd4 was successfully assayed in the TPP dataset. However, significant thermal shifts for Nedd4 were only observed at low NAB2 concentrations in our previous work and not at the 20 µM NAB2 concentration used in the current work.^29^ The absence of a T_m_ shift at higher ligand concentrations indicates that the binding reaction is more complex than a 1:1 binding event.^29^

### Rab1a as a putative, previously unrecognized target of NAB2

Rab1a, the protein identified as a consistent hit across the one-pot TPP and SPROX analyses, is an interesting target of NAB2, given this protein’s functional role in the cell. Rab1a, one of two isoforms of Rab1, is member of a family of small GTPases that regulate endomembrane trafficking and transport processes across the cell.^33,37,38^ Importantly, a number of these Rab-mediated pathways are known to be disrupted in PD and other neurodegenerative disorders.^37^ These GTPase proteins act as molecular switches and regulate trafficking by adopting functionally distinct conformations in a GTP/GDP-dependent manner.^39^ At a molecular level, binding of a Rab protein to GTP induces a conformational switch to its “on” position, enabling interaction of the Rab/GTP complex with effector proteins that regulate key steps of vesicular trafficking (tethering, motility, fusion, etc.).^33,34,37,38,40,41^ The Rab signaling process is tightly regulated by GTPase Activating Proteins (GAPs) that stimulate hydrolysis of GTP to GDP, serving to switch the Rab “off”, and Guanine Exchange Factors (GEFs) that induce exchange of GDP for GTP, switching the Rab back “on” (**Figure 3D**).

Rab1 has been established as a regulator of trafficking from the endoplasmic reticulum (ER) to the Golgi apparatus,^35,36,42–44^ a process that is disrupted by α-synuclein toxicity^35,36,45–47^ and restored by NAB2 treatment in models of PD (**Figure 3E**).^6,7^ It has been experimentally demonstrated that levels of Rab1 are decreased upon induction of α-synuclein toxicity,^48^ while Rab1 overexpression rescues cells by alleviating aggregate-induced toxicity.^36,47^ This effect is present not only in models of α-synuclein toxicity but in models of ALS, where Rab1 overexpression can restore defects associated with TDP-43, FUS or SOD1-related toxicity in cellular models of ALS.^37,42,43,49,50^ This makes the involvement of Rab1 in NAB2’s mechanism of action even more interesting as the NAB scaffold was first found to have phenotypic activity in models of ALS (TDP-43 toxicity) prior to advancement in models of α-synuclein-associated PD.^6^ These previous investigations indicate that the utility of Rab1 and NAB2 as a target and lead could extend beyond PD.

Based on the phenotypic activity of NAB2 and our knowledge of Rab1-mediated ER-to-Golgi trafficking, we hypothesize that NAB2 targets Rab1 and serves to stimulate downstream Rab1-mediated trafficking. Mechanistically, this would be consistent with either stabilizing the GTP-bound “on” state of Rab1 or by destabilizing the GDP-bound “off” state of Rab1. While this could occur by affecting the conformational properties of Rab1 directly, it could also occur through disruption of regulatory protein-protein interactions that serve to maintain Rab1 in its “off” state. These hypotheses are reinforced by the results obtained in the initial one-pot TPP and SPROX analyses, where Rab1 was stabilized in SPROX and destabilized in TPP, potentially through disruption of a protein-protein interaction after a direct binding event between NAB2 and Rab1 and/or by NAB2 binding to Rab1 inducing a conformational change, the latter of which is supported by our *in vitro* studies below.

Interestingly, no other Rab proteins were identified as hits in the one-pot TPP/SPROX analysis despite the high degree of structural similarity across the Rab GTPase family.^34,51^ Both Rab1 isoforms (Rab1a and Rab1b) were successfully analyzed in the TPP experiment while only Rab1a was identified in the SPROX experiment. The potential of small molecule-mediated Rab1 stimulation to alleviate PD-associated toxicity is promising, but the prospect of selectively targeting Rab1, or any other Rab GTPases, with high selectivity and in a functionally distinct manner (i.e. stimulating “on” or “off” position preferentially) has posed a challenge.^40^ There are, to our knowledge, no reports of ligands selective for Rab1, and few reports of ligands that preferentially bind to other Rab GTPases with selectivity.^38,40,52,53^ Therefore, the identification of Rab1 only as a hit in this screen could indicate that NAB2 binding and selectivity to Rab1 may be driven by Rab1-specific protein-protein interactions that would differentiate Rab1 as a target from the other, structurally related GTPases.

To investigate the interaction of NAB2 and Rab1, the relationship between newly identified target Rab1 and previously established target Nedd4, and the mechanism of NAB2 and Rab1 in the rescue of α-synuclein toxicity, a series of biochemical analyses were pursued.

### Validation of NAB2-dependent hit Rab1a reveals NAB2 binding is selective for the Rab1a-GDP complex

Previous studies have shown that Rab1 conformation and subsequent regulatory protein-protein interactions are dictated by the identity of the nucleotide (GTP or GDP) to which Rab1 is bound.^34,38^ We hypothesized that NAB2 binding may be specific to a particular Rab1 conformation (i.e. Rab1-GTP or Rab1-GDP complex). To investigate Rab1a stability in a NAB2-and nucleotide-dependent manner, we first generated Rab1a as a purified, untagged recombinant protein. With recombinant Rab1a in hand, we sought to screen the stability of the protein in response to nucleotide binding and NAB2 treatment in a rapid but robust manner. To this end, a protein thermal shift assay (PTSA) was used to screen Rab1a thermostability in which temperature-dependent protein unfolding is indicated by an increase in signal of a fluorescence dye, SYPRO Orange, that exhibits increased fluorescence upon binding to hydrophobic residues that are exposed upon protein denaturation.^54^ Using the PTSA workflow, Rab1a stability was first screened in response to nucleotide binding. In this case, Rab1a was screened in its apo form and after pre-incubation with GDP or non-hydrolysable GTP analogue GppNHp (**Figure 4A**). This experiment revealed that Rab1a was significantly stabilized (indicated by positive increase in melting temperature, T_m_) in the presence of GDP or GppNHp relative to the apo control. It is known that the affinity of the GDP and GTP interactions with Rab proteins is high, with binding affinities of the nucleotides in the nanomolar range.^55^ It has also been established that GTP binds more tightly to Rab proteins than GDP. Consistent with this information, we observed a larger shift in T_m_ in the presence of GppNHp than in the presence of GDP, indicating that GppNHp binds with higher affinity than GDP.

**Figure 4:**
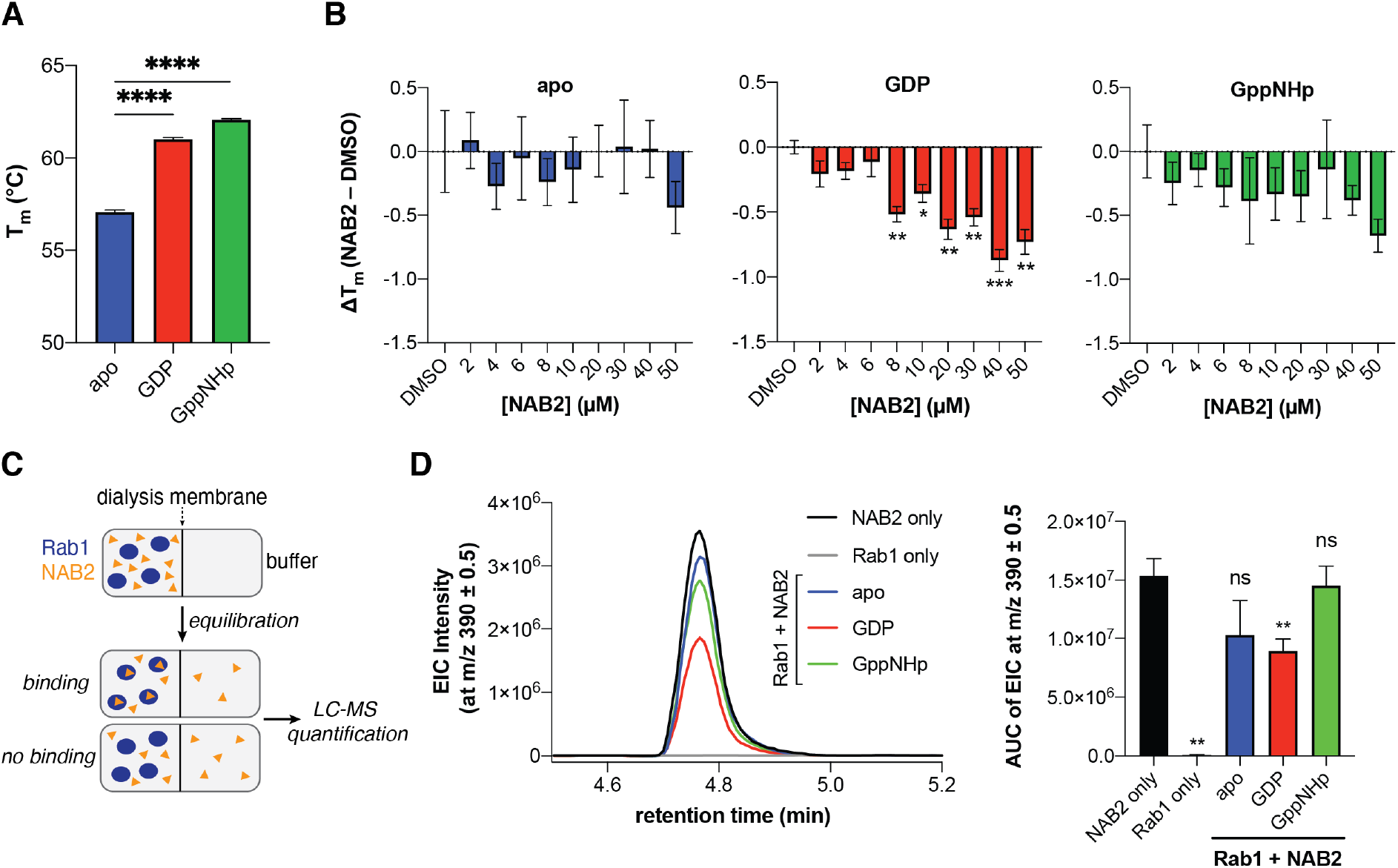
Protein thermal shift assay (PTSA) and equilibrium dialysis indicate NAB2 binds to Rab1a *in vitro* in a GDP-dependent manner. **(A)** Recombinant Rab1a is stabilized in the presence of GDP or non-hydrolysable GTP analogue GppNHp. **(B)** Recombinant Rab1a in its apo-, GDP-, and GppNHp-bound forms was treated with a concentration gradient of NAB2 and analyzed by PTSA, revealing nucleotide and [NAB2]-dependent shifts in Rab1a thermostability. Data shown as mean ± s.e.m. of triplicate measurements where ΔT_m_ was calculated as the difference of the mean of NAB2-treated sample relative to DMSO control for each Rab1a form and error was propagated appropriately. For **A** and **B**, data was collected on a Roche LightCycler 480 qPCR instrument and analyzed using Prism GraphPad. Significance was determined using a t-test with Welch’s correction for unequal variances where **** = p < 0.0001, *** = p < 0.001, ** = p < 0.01, * = p < 0.05). **(C)** NAB2 binding was confirmed by equilibrium dialysis. Recombinant Rab1a (apo-, GDP- or GppNHp-bound forms) was treated with NAB2 and loaded into a micro-dialysis chamber and allowed to equilibrate with the buffer blank for 24 h at 4 °C prior to LC-MS detection. All conditions were prepared in triplicate. **(D)** NAB2 in the buffer chamber was subsequently detected by LC-MS analysis and quantified by integration of the extracted ion chromatogram curve (EIC). Representative EIC curves for NAB2 only (positive control), Rab1 only (negative control), and Rab1 + NAB2 samples are shown *(left)*. Integration of EIC curves (AUC) for each condition are shown as mean ± s.d. of the triplicates *(right)*. Data was analyzed using Prism GraphPad and significance was determined using a t-test with Welch’s correction for unequal variances where ** = p < 0.01.

With this information in hand, the stability of Rab1a in its apo-, GDP-, and GppNHp-bound forms was subsequently screened in the presence of a NAB2 gradient (**Figure 4B**). This analysis revealed that Rab1a bound to GDP is more significantly destabilized in the presence of [NAB2] than in the presence of GppNHp. This result indicates that NAB2 binding is more selective for the GDP-bound form. Further, as there is no [NAB2]-dependent shift in the stability of apo-Rab1a, we infer that NAB2 binding is likely allosteric. If NAB2 binding was competitive, we anticipated that a T_m_ shift between apo and apo + NAB2 would be observed as it was with the initial nucleotide screen (**Figure 4A**) Therefore, the lack of [NAB2]-dependent shift in apo-Rab1a indicates an allosteric as opposed to a competitive binding mechanism where NAB2 binds in the enzymatic active site. Finally, the magnitude of the thermostability shift (ΔT_m_) induced by NAB2 binding to Rab1a-GDP is smaller than that induced by GDP or GppNHp to the apo form (< 1 °C compared to ∼3-5 °C shifts in T_m_, respectively). This suggests that NAB2 may exhibit a weaker binding affinity than the nucleotides themselves. This is consistent with the types of intermolecular forces that would drive ligand binding, where the nucleotides form ionic interactions through the di- or tri-phosphate group while NAB2 is not charged. Further, it is likely that an allosteric binding event would exhibit a lower affinity than binding in a discrete enzymatic active site. While the specific binding mode of NAB2 to Rab1-GDP has yet to be elucidated, this initial PTSA experiment indicates that NAB2 does affect Rab1 stability in a concentration- and nucleotide-dependent manner. Moreover, the consistency of these PTSA results with the purified Rab1a construct and the TPP results with Rab1a in the cell lysate suggest that the observed decrease in T_m_ upon NAB2 binding to Rab1a results from a conformational change in Rab1a upon NAB2 binding and not the disruption of Rab1a binding to other proteins, which would only be present in the TPP experiment.

To further validate the selectivity of NAB2 for Rab1a-GDP, we employed equilibrium dialysis with LC-MS detection to screen NAB2 binding to the Rab1a forms. In this experiment, Rab1a (apo, GDP- or GppNHp-bound) was treated with NAB2 and loaded into one chamber of a micro-dialysis cassette. The other chamber was loaded with buffer blank, and samples were allowed to equilibrate over 24 h at 4 °C followed by LC-MS analysis of samples from the buffer chamber of each cassette. In this experiment, we anticipated that a binding event would result in a lower effective concentration of NAB2 in the blank after equilibration relative to a positive control (NAB2 only) or a sample in which no binding occurred (**Figure 4C**). Effective NAB2 concentration was determined by integration of extracted ion chromatograms (EIC) at m/z 390.1 ([M+H]^+^ for NAB2) across three replicates of each experimental condition (**Figure 4D**). The EIC data reveals that the effective concentration of NAB2 is significantly lower only in the presence of Rab1a-GDP compared to positive control (NAB2 only), and that there is no statistically significant difference in NAB2 concentration in the Rab1a-apo and GppNHp-bound forms relative to the positive control (**Figure 4D, right**). This result provides an orthogonal and complementary demonstration that NAB2 binding to Rab1a is selective for the GDP-bound form of the enzyme.

### PTSA screening of NAB analogues reveals substituent-dependent effects on Rab1 destabilization

To expand our understanding of Rab1a as a potential target of NAB2, we sought to exploit previously established SAR analyses to look for structure-dependent alterations in NAB-induced shifts in Rab1a thermostability. To this end, the high-throughput PTSA platform was adapted for screening of a small library of NAB analogues that were previously identified to have various activities in the phenotypic rescue of α-synuclein toxicity (**Figure 7**).^6^ Each of these analogues, which have phenotypic activities ranging from EC_40_ = 4.5 to 32.75 µM or were included as phenotypically inactive controls (NAB17, NAB19), was incubated with recombinant Rab1a in its apo-, GDP- or GppNHp-bound form. The stability of Rab1a in the presence of each of these ligands was measured relative to DMSO controls, and stability shifts were calculated as ΔT_m_ (T_m_^NAB^ – T_m_^DMSO^). As in the analysis of NAB2 binding to Rab1a, larger ΔT_m_ values were observed in the GDP and GppNHp-bound forms relative to apo-Rab1a. Close analysis of the relationships between structural modification and ΔT_m_ reveals some information about the substituents that affect the NAB-dependent shift in Rab1a stability. For instance, for Rab1a-GDP, no shift in ΔT_m_ is observed in the presence of NAB17, a phenotypically inactive derivative that lacks the benzdiimidazole ring closure. There are minor changes induced by alteration of methyl substituents on the *N*-aryl moiety, as evidenced by ΔT_m_ in the presence of NAB2 versus NAB1 (*m-*methyl), NAB13 (no methyl substituents), and NAB19 (*p-*methyl). Meanwhile, alteration of the 2-chloro substituent on the benzylamine region of the lead NAB scaffold (NAB1) to 2-methoxy (NAB9) or 2-fluoro (NAB15) induces little change and a small but not statistically significant change in ΔT_m_, respectively. The trends are more pronounced in the Rab1a-GppNHp samples, indicated by clear shifts in ΔT_m_ induced by substitutions in the C(2) position of the benzylamine moiety (NAB1, 2-chloro; NAB9, 2-methoxy; NAB15, 2-fluoro). Variation of substituents on the *N-*aryl moiety also affect ΔT_m_, where neither NAB1 (*m-*methyl) and NAB13 (no methyl substituents) induced significant ΔT_m_ values relative to DMSO controls (ΔT_m_ ∼ 0 °C). However, NAB2 (2,5-dimethyl), NAB4 (*m-*ethyl) and NAB19 (*p-*methyl) induced similar changes in ΔT_m_.

**Figure 5:**
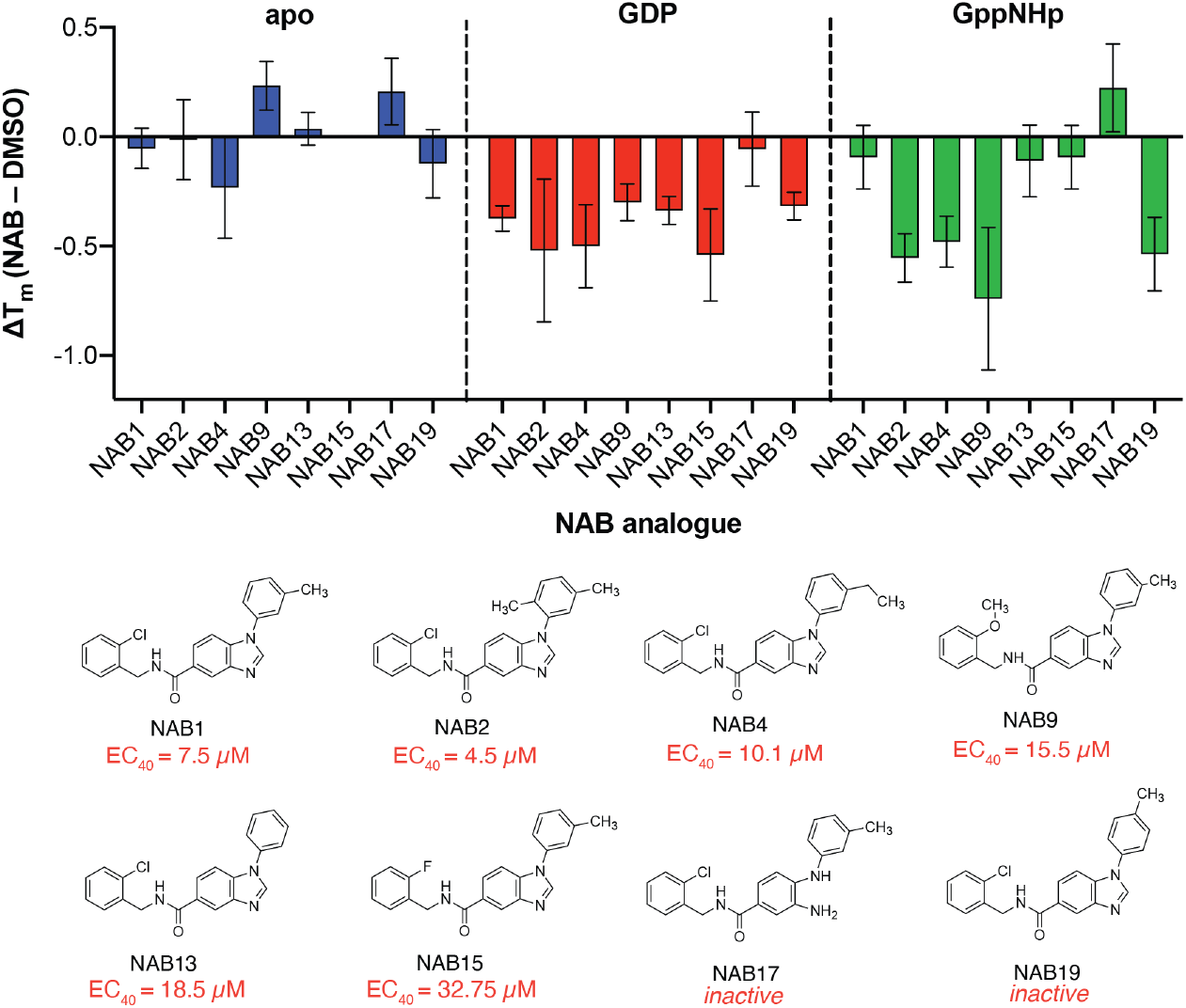
Variation of substituents on the NAB scaffold affects ΔT_m_ of recombinant Rab1a in a nucleotide-dependent manner but does not correlate with previously reported phenotypic activity. Recombinant Rab1a (2 µM) stability was screened after incubation with eight different NAB analogues (50 µM) via PTSA analysis. Shifts in melting temperature show no clear correlation with phenotypic activity (as determined in yeast-based phenotypic screening reported by Tardiff and co-workers)^6^, but trends in substituent effects in GDP- and GppNHp-bound Rab1a are revealed. Data shown as mean ± s.d. of triplicate measurements where ΔT_m_ was calculated as the difference of the mean of NAB-treated sample relative to DMSO control for each Rab1a form and error was propagated appropriately. Data collected on Roche LightCycler 480 qPCR instrument and analyzed using Prism GraphPad.

**Figure 6:**
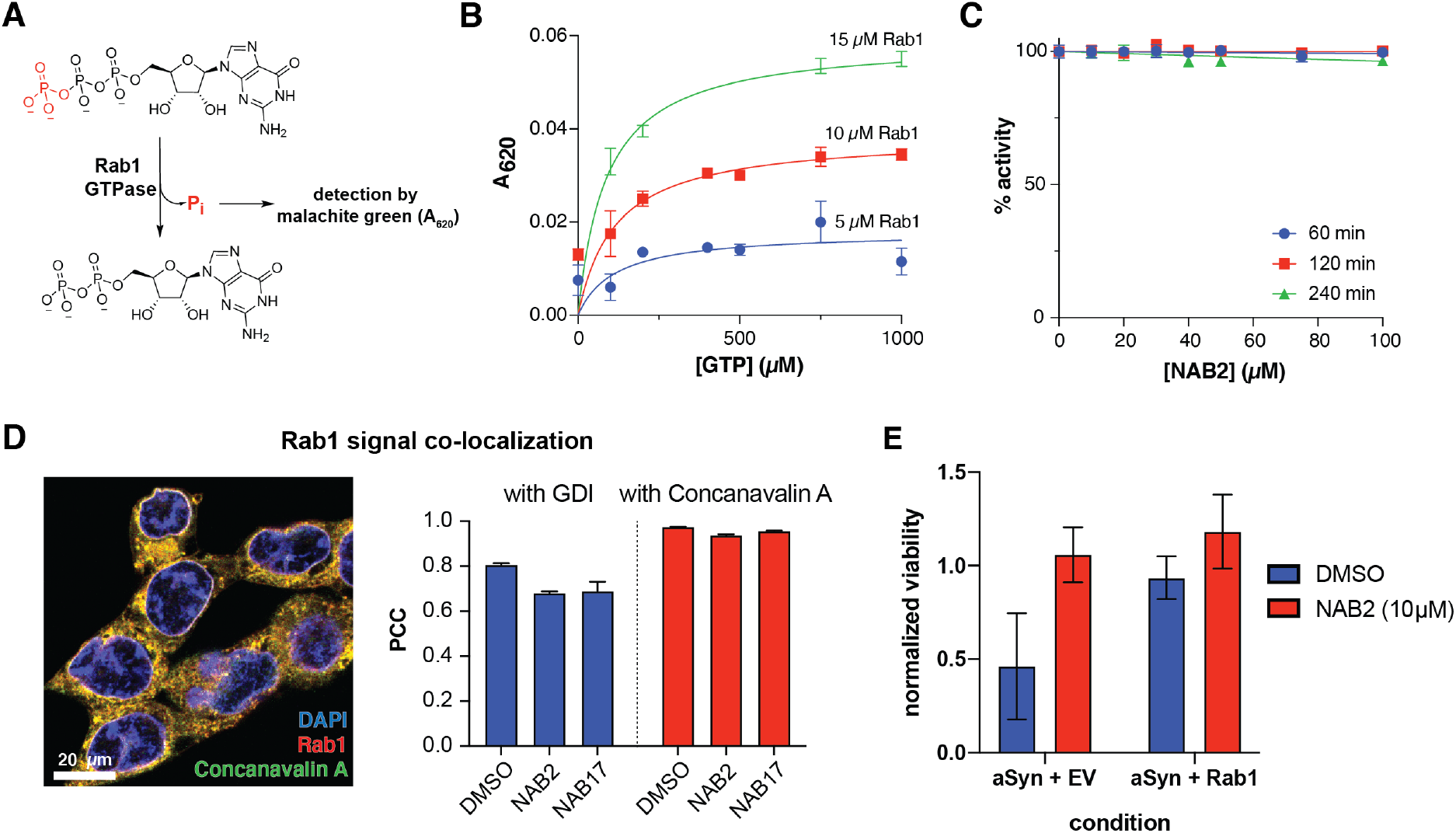
Rab1a overexpression and NAB2 treatment improve viability of α-synuclein toxic cells but NAB2 does not alter Rab1 GTPase activity or localization. **(A)** Rab1a exhibits GTPase activity. As GTP is enzymatically hydrolyzed to GDP, free orthophosphate is released as a product of the reaction. Free orthophosphate can be detected in a colorimetric manner through formation of a complex with malachite green and molybdenum. **(B)** Orthophosphate formation, measured by malachite green complex formation and absorbance at 620 nm, occurs in a [Rab1]- and [GTP]-dependent manner. **(C)** Rab1 activity is not affected by NAB2 treatment (quantified as % activity relative to DMSO controls). Data in **B** and **C** shown as mean ± s.e.m. of triplicate measurements. **(D)** Representative merged immunofluorescence microscopy image is shown for monitoring the co-localization of Rab1 with membrane marker Concanavalin A. Quantification of protein co-localization was measured as Pearson correlation coefficient (PCC) and indicates that NAB2 treatment does not alter Rab1a co-localization with GDI or Concanavalin A relative to NAB17 control. Co-localization was measured volumetrically (across the z-stack image) for a minimum of fifteen cells across three microscopy frames. Data presented as mean ± s.e.m. of PCC from all individual cells measured. Images collected with Zeiss Airyscan 880 confocal microscope and analyzed with Imaris. Representative images prepared with ImageJ,^64^ and data analysis was performed with Prism GraphPad. **(E)** NAB2 treatment increased viability in α-synuclein toxic cells relative to DMSO controls (left), an effect that was phenocopied by co-expression of α-synuclein with Rab1a (right). NAB2 treatment induces a slight increase in viability of α-synuclein toxic cells overexpressing Rab1a. Data shown as mean ± s.e.m. of four biological replicates prepared and analyzed in parallel.

**Figure 7:**
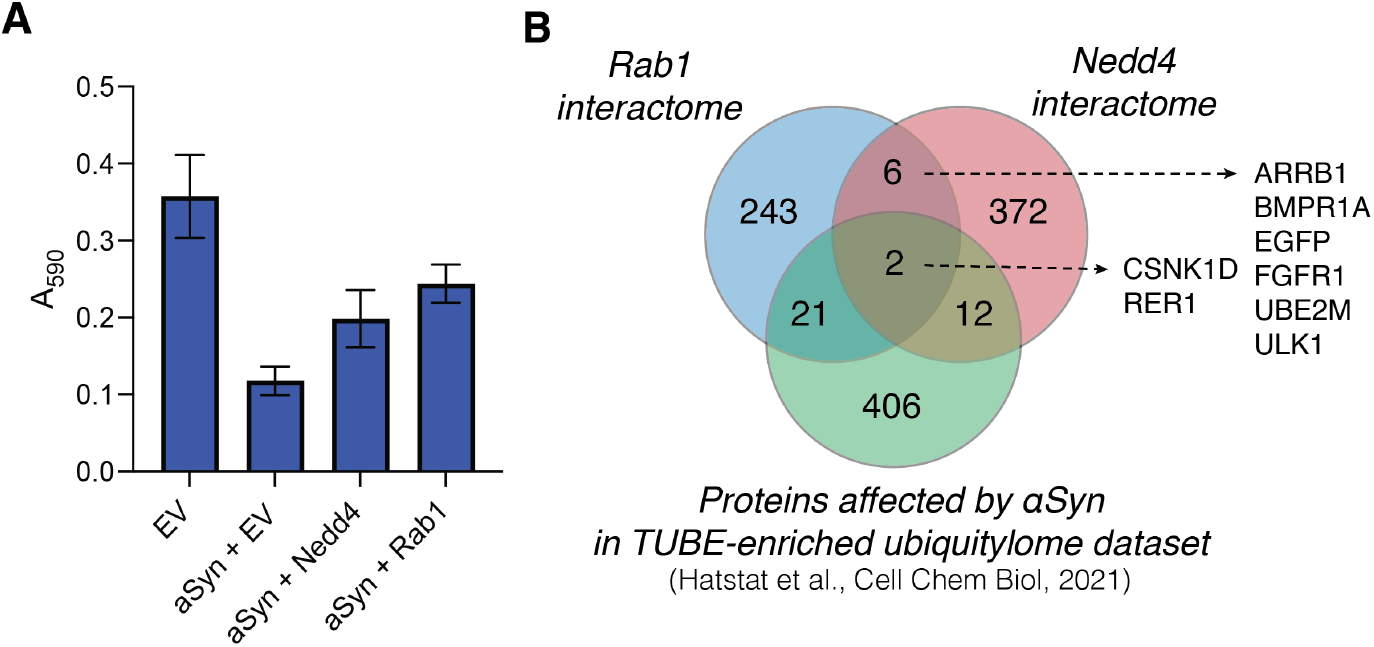
Preliminary exploration of link between Nedd4 and Rab1a in the NAB2-dependent rescue of α-synuclein toxicity. **(A)** Rab1a and Nedd4 overexpression both independently improve cell viability relative to α-synuclein overexpression alone. Viability of SHSY5Y cells was monitored at 48 hours post-transfection via MTT assay in the presence of α-synuclein overexpression (as a model of toxicity) with co-expression of empty vector (EV) or with putative NAB2 targets Nedd4 and Rab1a. Data shown as mean ± s.e.m. of four biological replicates prepared and analyzed in parallel. **(B)** Cross-reference of Nedd4 and Rab1a interactomes (retrieved from BioGrid database) with proteins previously identified in the α-synuclein toxicity-dependent ubiquitylome reveals a small pool of functionally diverse regulatory proteins that interact with both proteins of interest and are affected by α-synuclein toxicity.

The differential trends in ΔT_m_ in response to structural variations of the NAB scaffold between the Rab1a-GDP and Rab1a-GppNHp forms suggest that NAB binding is affected by a nucleotide-dependent change in Rab1a conformation and indicate that binding may occur in the region of the protein that is involved in its conformational “molecular switch” activity. While this experiment provides some insight into the structural features of the NAB scaffold that could contribute to Rab1a binding, it reveals that there is not a clear correlation between observed phenotypic activity and binding affinity (as inferred by magnitude of ΔT_m_). This result suggests that NAB phenotypic activity may be affected by more than just the affinity for Rab1a alone.

### NAB2 does not alter Rab1a activity or localization in vitro

To begin investigating the effect of NAB2 on Rab1a and associated trafficking, we first sought to characterize the effect of Rab1a activity in response to NAB2 treatment. Rab1a, as a GTPase, cleaves GTP to GDP via enzymatic hydrolysis of a phosphorous-oxygen bond in the triphosphate tail of GTP.^38,55^ This reaction produces free orthophosphate as a side product, thus making the Rab1 enzyme amenable to use in assays where orthophosphate formation is monitored in an enzyme-dependent manner. In this case, we opted to employ a standard malachite green assay where orthophosphate is detected by complex formation between inorganic phosphate, molybdenum and malachite green (**Figure 6A**).^56^ Complexation of these assay components generates a colorimetric signal between 600-660 nm proportional to the levels of free orthophosphate. While this assay must be conducted in a discontinuous format, it provides sufficient sensitivity for a preliminary screen of Rab1a activity in response to NAB2 treatment. To begin, we validated the assay by measuring A_620_ in a [Rab1a]-and [GTP]-dependent manner (**Figure 6B**). The A_620_ signal as a measure of orthophosphate formation is proportionate to [Rab1a], and all reactions demonstrate [GTP]-dependent activity until saturation is achieved at V_max_. It should be noted that intrinsic activity of Rab GTPases is very slow, and that activity *in cellulo* is stimulated by GTPase Activating Proteins (GAPs) (**Figure 3D**).^57^ As this assay was conducted with recombinant Rab1 alone, assay reactions were conducted for an extended 3 hour duration to provide sufficient time for detectable hydrolysis to occur. This analysis confirms that the recombinant Rab1 enzyme exhibits the expected activity.

With the assay platform established and validated, Rab1a activity was next measured in the presence of NAB2 in a time- and concentration-dependent manner. Rab1a (5 µM) was incubated with 1 mM GTP in the presence of 0–100 µM NAB2 for 60, 120, and 240 minutes. As NAB2 binding to Rab1a was shown to be selective for the GDP-bound form, we anticipated that NAB2 would not alter the enzymatic activity of Rab1a. In all cases, no change in Rab1-dependent GTP hydrolysis was observed relative to vehicle treated controls (0 µM NAB2) (**Figure 6C**). This result furthers our hypothesis that the role of Rab1a in NAB2-mediated alleviation of α-synuclein toxicity and restoration of ER-to-Golgi trafficking occurs at a regulatory level as opposed to through changes in the intrinsic activity of Rab1. Therefore, further analyses of Rab1a regulation were explored in a NAB2-dependent manner.

As with all members of the Rab GTPase family, Rab1a regulation of endomembrane trafficking is dependent upon geranylgeranylation for lipid-mediated anchoring to membranes.^55,58,59^ This localization is further regulated by guanosine nucleotide dissociation inhibitor protein (GDI), which can sequester prenylated Rab1a away from the membrane by binding to the lipid-modified enzyme (**Figure 3D**).^60,61^ To determine if NAB2 treatment stimulates trafficking by increasing the resonance time of Rab1a at the membrane (thus enabling Rab1a to exist in its membrane-associated, GTP-bound “on” state for a longer time), the localization of Rab1a was monitored in a NAB-dependent manner. In these experiments, which included analysis by both subcellular fractionation and confocal microscopy-based measurement of co- localization, SHSY5Y cells were treated with NAB2, phenotypically inactive derivative NAB17, or DMSO vehicle control. Subcellular fractionation experiments reveal that Rab1a is primarily present in the membrane bound fraction, and NAB2 treatment did not induce a significant alteration in Rab1a localization relative to NAB17 control (**Figure S1**). Rab1a localization was further monitored by immunofluorescence-coupled confocal microscopy where the degree of Rab1a co-localization with GDI (as a measure of sequestration to the cytoplasm) and membrane marker Concanavalin A_62_ (as a measure of localization to endomembrane organelles) was measured (**Figure 6D**). As in co-localization experiments previously reported,^29^ co-localization was determined by quantifying the signal for each immunofluorescent probe volumetrically (across all slices of a z-stack) in a minimum of fifteen cells across three microscopy frames. Signal co-localization was measured quantitatively, where the intensity of Rab1a signal and marker signal in each pixel of the quantified area was correlated and measured as Pearson’s correlation coefficient (PCC).^63^ In this analysis, a positive correlation value (0 < PCC < 1) indicates a positive degree of correlation and a negative value (–1 < PCC < 0) indicates inverse localization. Comparison of PCC values across experimental conditions reveals that NAB2 treatment does not alter Rab1a co-localization with GDI or Concanavalin A relative to NAB17 controls. There is a slight decrease in co-localization with GDI and Concanavalin A in NAB2-treated samples relative to DMSO control. Despite this, since NAB17 is confirmed to be a phenotypically inactive derivative, the lack of difference between PCC values for NAB2 samples relative to NAB17 controls indicates that the NAB2 mechanism is independent of changes in Rab1a co-localization.

### NAB2 treatment phenocopies Rab1 overexpression

It has been previously determined that α-synuclein toxicity disrupts Rab homeostasis^48^ and induces trafficking defects in the endomembrane system. Specifically, it has been shown that α-synuclein toxicity decreases levels of Rab1 and that Rab1 overexpression restores trafficking processes and minimizes toxicity phenotypes induced by α-synuclein in both cellular and animal models.^35,36,65^ Since Rab1 has not been previously studied in the context of NAB2-mediated alleviation of α-synuclein toxicity, we sought to compare the effect of Rab1 overexpression with that of NAB2 treatment. To this end, a cell-based assay was established to monitor the viability of neuroblastoma-derived SHSY5Y cells in the presence of α-synuclein toxicity and the ability of putative target Rab1a to rescue viability. SHSY5Y cells were transfected with a plasmid for constitutive expression of α-synuclein in combination with an empty vector control or a Rab1a expression plasmid. The viability of cell populations was subsequently measured at 24 or 48 hours post transfection relative to empty vector (EV) controls using a 3-(4,5-dimethylthiazol-2-yl)-2,5- diphenyl tetrazolium bromide (MTT) viability assay.^66^ In the MTT assay, cell viability is measured as a function of cellular metabolic activity. Viable cells convert MTT into formazan through activity of metabolic NADPH-dependent oxidoreductase enzymes, enabling colorimetric analysis of cell viability (measured as absorbance at 590 nm). Using this platform, the viability of cells in the presence of α-synuclein or α-synuclein co-expressed with Rab1a (**Figure 6E**). In this first analysis, MTT-based viability measurements indicate that SHSY5Y cell viability is decreased upon expression of α-synuclein relative to EV control, but the viability is partially restored when α-synuclein is co-expressed with Rab1a.

To determine if NAB2 treatment phenocopies Rab1a overexpression, the effect of NAB2 treatment on cell viability in the presence of α-synuclein alone or co-expressed with Rab1a was measured (**Figure 6E**). This analysis revealed that NAB2 treatment improved viability in the presence of α-synuclein overexpression alone, and the degree of viability restoration was similar to that of Rab1a co-expression. Further, there is a slight but not significant increase in viability of cells co-expressing α-synuclein with Rab1a and also treated with NAB2. This analysis indicates that NAB2 treatment phenocopies Rab1a overexpression and further supports our hypothesis that NAB2 treatment is stimulating the “on” state of Rab1a and improving downstream trafficking as a means of combatting α-synuclein toxicity. While the nature of Rab1a stimulation is still not fully clear, the evidence collected to date supports our initial hypotheses about the role of NAB2 and Rab1a in the rescue of α-synuclein toxicity.

### Putative links between Nedd4 and Rab1a in the NAB2 mechanism of action

To expand our understanding of the mechanism of action of NAB2 in the rescue of α- synuclein toxicity, we conducted a preliminary screen of the role of Rab1 and Nedd4 in the restoration of α-synuclein toxicity. To this end, the SHSY5Y-based MTT viability assay was conducted in which α-synuclein was overexpressed via co-transfection with empty vector control or plasmids for expression of Rab1a or Nedd4 (**Figure 7A**). Following transfection and protein expression, cell viability was measured by MTT reduction and revealed that α-synuclein overexpression decreased viability relative to empty vector controls while both Rab1a and Nedd4 overexpression improved cell viability relative to expression of α-synuclein alone. This provides further evidence that both Nedd4 and Rab1 may be implicated in the rescue of α-synuclein toxicity.

We next sought to investigate any potential links between Nedd4, the target initially identified through chemical genetic screening, and Rab1a, the protein identified in our chemoproteomic analysis of NAB2 targets. As Rab1a has not been previously annotated as a Nedd4 substrate (as indicated in BioGrid interactome database)^67,68^ and Rab1a lacks a canonical PY motif for recognition by Nedd4, we anticipate that Rab1a is not a direct substrate of Nedd4. Instead, we hypothesize that Rab1a and Nedd4 could be independent targets of NAB2 or that Nedd4 could regulate Rab1a activity indirectly, such as through modulating the state or abundances of Rab1a regulators or interactors through ubiquitination-induced degradation. To being to investigate the latter, we sought to identify shared pathways or functions of the two enzymes. The experimentally annotated interactomes were retrieved from the BioGrid^67,68^ database and cross-referenced (**Figure 7B**).^69^ To further explore these interactomes in the context of α-synuclein toxicity and ubiquitin signaling, we also cross-referenced both interactomes with the set of proteins significantly altered in the ubiquitylome following induction of α-synuclein toxicity that we previously reported.^29^

Through this analysis, a small group of proteins is revealed that have been shown to interact with both Rab1a and with Nedd4 in separate experimental analyses (**Figure 7B**). The majority of the proteins identified as shared interactors exhibit regulatory functions in the cell where they participate in signal transduction or regulation (as determined through Gene Ontology annotation).^70,71^ For example, EGFR and FGFR1 are transmembrane signal receptors while ULK1 is a kinase involved in autophagy. ARRB1 (beta-arrestin-1) is a scaffolding protein involved in transducing signals from G-protein coupled receptors (GPCRs). Cross-reference of the Nedd4 and Rab1a interactomes with our α-synuclein-dependent ubiquitylome dataset reveals two proteins that are present in all three datasets: CSNK1D and RER1. Both of these proteins are involved in endomembrane trafficking where CSNK1D regulates vesicle budding, membrane invagination, and endocytosis^72,73^ while RER1 is a transmembrane protein localized to the Golgi membrane that regulates vesicle mediated transport between the ER and Golgi.^74–76^ Further, both proteins have been directly studied in the context of α-synuclein toxicity or related neurodegenerative trafficking defects outside of our proteomic evaluation.^77–80^ The identification of CSNK1D and RER1 as putative links between Nedd4, Rab1a and α-synuclein toxicity is particularly exciting in our exploration of NAB2-mediated rescue of α-synuclein toxicity and we anticipate that the relationships of these enzymes will be explored further in future analyses.

## Conclusion

Collectively, these data suggest that Rab1a is a previously unrecognized protein affected by NAB2 treatment in the rescue of α-synuclein toxicity. The role of Rab1a in regulation of ER-to-Golgi trafficking, the process disrupted by α-synuclein toxicity and restored by NAB2 treatment, has been thoroughly and robustly established in various experimental models. ^4,6,42–47,49,50,65^ These prior studies, in combination with data from this work, suggests that that Rab1 is likely a member of the protein network implicated in the NAB2 mechanism of action. We discovered that NAB2 binds to Rab1-GDP selectively over the apo- or GppNHp-bound forms of the protein, and NAB2 treatment phenocopies Rab1a overexpression in the restoration of cell viability without altering Rab1a activity or localization. Based on these results, we hypothesize that NAB2 binding to Rab1-GDP induces a shift in Rab1 regulation and stimulates a “gain-of-function” effect to restore or preserve control of normative trafficking processes.

## Acknowledgements

The authors would like to thank the Duke University Department of Chemistry Shared Instrumentation Facility Director, Dr. Peter Silinski, for supporting the mass spectrometry analyses in this work. The authors also acknowledge Lydia Stariha for assistance in development of LC-MS method for equilibrium dialysis experiments. Finally, the authors thank the members of the McCafferty lab for their thoughtful feedback on the project and manuscript.

