## Supplemental Materials for "Chemoproteomic-enabled characterization of small GTPase Rab1a as a target of an *N*-arylbenzdiimidazole ligand’s rescue of Parkinson’s-associated cell toxicity"

### Methods

#### Expression and purification of recombinant Rab1a

Purification of Rab1a was adapted from previously reported methods.[1] Briefly, pGEX-4T-Rab1a was obtained from Addgene (#49565) and modified by site-directed mutagenesis for inclusion of a TEV protease cut site downstream of the N-terminal GST tag to allow for facile separation of the GST affinity tag from Rab1a. Primers were designed using NEBaseChanger and synthesized by Eton Biosciences as follows: 5'-TTTTCAGGGCACCATGTCCAGCATGAATCC-3' (FWD) and 5'-TACAGGTTTTCGGATCCACGCGGAACCAC-3' (REV). Primers were resuspended from lyophilized stocks and used for Q5 mutagenesis according to manufacturer protocols.

Following confirmation of successful mutagenesis by sequencing (Eton Biosciences), pGEX-4T-Rab1a was transformed into BL21(DE3)-CodonPlus-RIL competent *E. coli*, and successful transformants were selected for by growth on an ampicillin selection plate. For expression, a starter culture (100 mL LB media) was inoculated from a streak of colonies on the selection plate and allowed to reach saturation by overnight growth at 37 °C. Expression cultures were inoculated from saturated starter culture (10 mL starter culture per 1 L LB media) and grown at 37 °C until $OD_{600}$ = 0.6 at which time expression was induced with the addition of IPTG to a final concentration of 100 µM and grown overnight at 21 °C. After growth, cells were harvested and pelleted for storage at –20 °C until use for purification.

For purification, the cell pellet was thawed on ice and subsequent resuspended in lysis buffer (20 mM Tris, 150 mM NaCl, pH 7.4 with 1X protease inhibitor cocktail, 40 µM PMSF and lysozyme). The cells were lysed, and the cell debris was removed from the lysate by ultracentrifugation. GST-Rab1a was purified from the supernatant of the lysate as follows: the supernatant was loaded onto a Glutathione Agarose column (Genesee Scientific) that was pre-equilibrated with base buffer (20 mM Tris, 150 mM NaCl, pH 7.4). The loaded column was washed with 10 column volumes of base buffer before elution with glutathione elution buffer (base buffer supplemented with 20 mM reduced glutathione). Eluant containing protein as indicated by absorbance on the UV trace at 280 nm was pooled into 3.5 kDa MWCO dialysis tubing. The dialysis sample was supplemented with TEV protease and the sample was allowed to dialyze overnight against 4 L dialysis buffer (base buffer supplemented with 1 mM β-mercaptoethanol). Following dialysis, dialysate was loaded onto a $Ni^{2+}$-charged immobilized metal affinity column (equilibrated with 20 mM Tris, 150 mM NaCl, pH 7.5). The column was subsequently washed with ten column volumes of base buffer followed by gradient elution (base buffer supplemented with imidazole, 0 to 500 mM gradient). Rab1a containing fractions were confirmed by SDS-PAGE analysis and were pooled for subsequent concentration and buffer exchange into base buffer followed by aliquoting for storage at 4 °C, –20 °C (in 40% glycerol), and –80 °C.

#### Synthesis of NAB analogues

Derivatives of the *N*-arylbenzdiimidazole scaffold were synthesized according to previously reported procedures as established by Tardiff and co-workers[2] and as described below. All characterization data of the NAB derivatives prepared for the studies discussed are consistent with previously reported characterization.

*Materials*: Unless otherwise stated, all reagents were purchased from either Sigma-Aldrich or Oakwood Chemicals and were used as received. $CDCl_3$ was purchased from Cambridge Isotope Laboratories and used as received.

*Characterization*: NMR spectra were obtained using Bruker Advance Neo - 500 MHz multinuclear spectrometer (Duke Chemistry NMR Facility) operating at 500 MHz for $^{1}H$ NMR and 126 MHz for $^{13}C$ NMR and are reported as chemical shifts (δ) in parts per million (ppm). Spectra were referenced internally according to residual solvent signals ($^{1}H$, $CDCl_3$ 7.26 ppm; $^{13}C$, $CDCl_3$ 77.0 ppm). Data for NMR spectra use the following abbreviations to describe multiplicity: s, singlet; br s, broad singlet; d, doublet; t, triplet; dd, doublet of doublets; td, triplet of doublets; tt, triplet of triplets; ddd, doublet of doublets of doublets; dddd, doublet of doublets of doublets of doublets; ddt, doublet of doublets of triplets; app dd, apparent doublet of doublets; m, multiplet. Coupling constants (J) are reported in units of hertz (Hz). High-resolution mass spectra (HRMS, m/z) were recorded on an Agilent 6224 LC/MS-TOF spectrometer using electrospray ionization (ESI, Duke University Department of Chemistry Shared Instrumentation Facility).

*General procedures for preparation of N-arylbenzdiimidazole (NAB) derivatives*: NAB derivatives were prepared by one of two routes wherein amide bond formation was followed by nucleophilic aromatic substitution ($S_NAr$) and heterocycle formation. Alternatively, derivatives were prepared by sequential $S_NAr$, amide bond linkage and heterocycle formation. The general procedures are as follows:

<u>*General Procedure A:*</u>

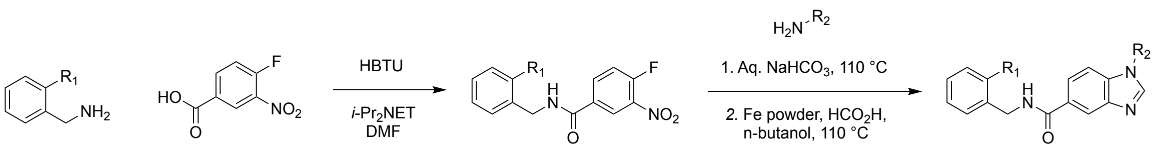

A 2-substituted benzylamine (1.1 equiv.) was added slowly to a stirred solution of 4-fluoro-3-nitrobenzoic acid (1 equiv.), HBTU (1 equiv.), *N,N*-diisopropylethylamine (1.1 equiv.), and DMF (2 mL/mmol of 4-fluoro-3-nitrobenzoic acid) at room temperature. After 30 min, the solution was diluted with ethyl acetate (5X DMF volume) and washed sequentially with water, 1M HCl (aq) (3X), 1M KOH (aq) (3X), and brine. The organic layer was dried over sodium sulfate, filtered, and concentrated under reduced pressure using a rotary evaporator. Purification of the residue by silica gel chromatography (10-45% EtOAc/Hexanes). Purified benzamide intermediate (1 equiv.) was subsequently added to a screw-top test tube containing sodium bicarbonate (2 equiv.), aniline (or substituted aniline derivative) (2 equiv.), and water (2 mL/mmol benzamide) was sealed with a Teflon screw cap, placed in a preheated oil bath at 110 °C, and stirred for 24 h. The reaction was cooled to room temperature and the mixture was poured onto ethyl acetate (sufficient volume for all solid to be dissolved). The solution washed sequentially with 1M HCl (aq) (3X), water (3X), and brine (3X). The organic layer was dried over sodium sulfate, filtered, and concentrated using reduced pressure via a rotary evaporator. The crude nitroaniline was subsequently dissolved in n-butanol (1.0 mL/mmol benzamide, 0.96 mL), transferred to a second screw-top test tube, and formic acid (1.0 mL/mmol benzamide, 0.96 mL), iron powder (9.6 mmol, 536 mg), and concentrated HCl (0.1 mL/mmol benzamide, 96 $\mu$L) were added to the solution. The test tube was sealed with a Teflon screw cap and stirred for 1 h at 110 °C. After cooling to room temperature, the mixture was poured onto a mixture of ethyl acetate and saturated $NaHCO_3$ (1:6 ratio) in a separatory funnel. The mixture was shaken with venting and solid sodium bicarbonate added until pH~12. The layers were separated, and the organic layer was washed with water (3X), dried and concentrated under reduced pressure. The residue was purified by silica gel chromatography using 40-100% EtOAc/Hexanes.

*General Procedure B:*

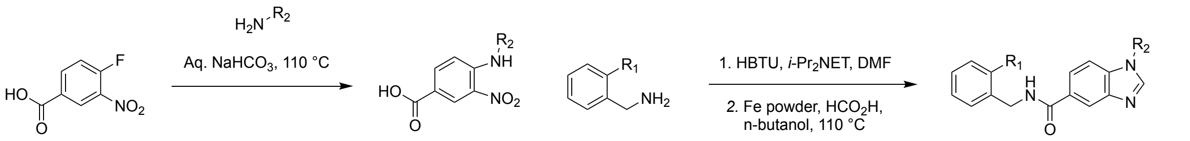

4-Fluoro-3-nitrobenzoic acid (1.0 equiv.) was reacted with an aniline derivative (1.0 equiv.) in the presence of $NaHCO_3$ (2.4 equiv.) and water (1.0 mL/mmol of 4-fluoro-3-nitrobenzoic acid) in a twist cap Teflon tube. The reaction mixture was heated to 110 °C and allowed to proceed for 24 h. Upon cooling to room temperature, the reaction mixture was transferred into a separatory funnel containing a 1:1 mixture of 1M HCl and EtOAc. The mixture was shaken vigorously, then the aqueous layer was separated and extracted twice with EtOAc. Organic layers were combined, dried with $Na_2SO_4$, filtered, and concentrated under reduced pressure. The crude nitroaniline was purified by flash chromatography using 10-30% EtOAc/Hexanes.

The nitroaniline intermediate (1.0 equiv.) was combined with HBTU (1.0 equiv.), *N,N*-diisopropylethylamine (1.2 equiv.) and DMF (2 mL/mmol 4-fluoro-3-nitrobenzoic acid used) and stirred. Benzylamine (1.0-2.0 equiv.) was added slowly. The reaction was allowed to proceed for 12 h before dilution with ethyl acetate and subsequent washing with water, 1M HCl (aq), 1M KOH (aq), and brine. The organic layer was dried over $Na_2SO_4$, filtered and concentrated under reduced pressure. The crude nitroaniline was subsequently dissolved in n-butanol (1.0 mL/mmol benzamide, 0.96 mL), transferred to a second screw-top test tube, and formic acid (1.0 mL/mmol benzamide, 0.96 mL), iron powder (9.6 mmol, 536 mg), and concentrated HCl (0.1 mL/mmol benzamide, 96 $\mu$L) were added to the solution. The test tube was sealed with a Teflon screw cap and stirred for 1 h at 110 °C. After cooling to rt, the mixture was poured onto a mixture of ethyl acetate and saturated $NaHCO_3$ (1:6 ratio) in a separatory funnel. The mixture was shaken with venting and solid sodium bicarbonate added until pH~12. The layers were separated, and the organic layer was washed with water (3X), dried and concentrated under reduced pressure. The residue was purified by silica gel chromatography using 40-100% EtOAC/Hexanes.

*Characterized products*

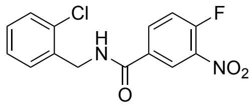

*N-(2-chlorobenzyl)-4-fluoro-3-nitrobenzamide.* 2-Chlorobenzylamine (0.71 mL, 6.0 mmol) was added slowly to a stirred solution of 4-fluoro-3-nitrobenzoic acid (925 mg, 5.0 mmol), HBTU (1.90 g, 5.0 mmol), *N,N*-diisopropylethylamine (1.05 mL, 6.0 mmol), and DMF (10 mL) at room temperature. After 30 min, the solution was diluted with ethyl acetate (50 mL) and washed sequentially with water, 1M HCl (aq) (3X), 1M KOH (aq) (3X), and brine. The organic layer was dried over sodium sulfate, filtered, and concentrated under reduced pressure using a rotary evaporator. Purification of the residue by silica gel chromatography (10-45% EtOAC/Hexanes; material loaded using toluene) provided the title compound as a yellow solid (0.77 g, 50% yield). $^{1}$H NMR (500 MHz, $CDCl_3$) δ 8.46 (dd, *J* = 7.0, 2.3 Hz, 1H), 8.11 (ddd, *J* = 8.8, 4.1, 2.3 Hz, 1H), 7.50 – 7.34 (m, 4H), 7.28 (m, 2H), 6.60 (s, 1H), and 4.75 ppm (d, *J* = 5.9 Hz, 2H). $^{13}$C NMR (126 MHz, $CDCl_3$) δ 164.30, 157.24 (d, *J* = 270.9), 134.96, 134.67 (d, *J* = 8.8), 133.73, 131.18 (d, *J* = 3.8) 130.28, 129.72, 129.33, 127.25, 125.2, 118.95 (d, *J* = 21.4), and 42.43 ppm. HRMS (ESI-TOF) m/z calcd for $C_{14}H_{10}ClFN_2O_3$ $[M+H]^+$ 309.0437, found 309.0445.

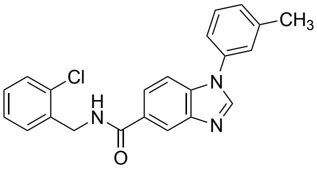

*N-(2-chlorobenzyl)-1-(m-tolyl)-1H-benzo[d]imidazole-5-carboxamide (NAB1).* NAB1 was prepared according to General Procedure A using 2-chlorobenzylamine to generate intermediate *N*-(2-chlorobenzyl)-4-fluoro-3-nitrobenzamide. The benzamide intermediate was purified and carried forward and reacted with 2-methylaniline (*o*-toluidine) under $S_NAr$ conditions and subsequent heterocycle formation to afford NAB2 as a white solid (74% yield). $^{1}$H NMR (500 MHz, $CDCl_3$) δ 8.26 (s, 1H), 8.17 (s, 1H), 7.87 (dd, *J* = 8.6, 1.7 Hz, 1H), 7.57 (d, *J* = 8.5 Hz, 1H), 7.54–7.44 (m, 2H), 7.43– 7.37 (m, 1H), 7.33 – 7.22 (m, 5H), 6.73–6.65 (m, 1H), 4.79 (d, *J* = 6.0 Hz, 2H), and 2.48 ppm (s, 3H). $^{13}$C NMR (126 MHz, $CDCl_3$) δ 167.50, 144.11, 143.90, 140.50, 135.81, 135.78, 131.74, 130.50, 129.97, 129.62, 129.31, 129.26, 129.05, 127.21, 124.71, 123.35, 121.16, 119.26, 110.78, 42.25, and 21.44 ppm. HRMS (ESI-TOF) m/z calcd $C_{22}H_{18}ClN_3O$ for $[M+H]^+$ 376.1138, found 376.1213.

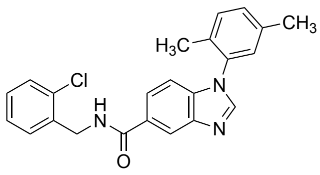

*N-(2-chlorobenzyl)-1-(2,5-dimethylphenyl)-1H-benzo[d]imidazole-5-carboxamide (NAB2).* NAB2 was prepared according to General Procedure A using 2-chlorobenzylamine to generate intermediate *N*-(2-chlorobenzyl)-4-fluoro-3-nitrobenzamide. The benzamide intermediate was purified and carried forward and reacted with 2,5-dimethylaniline under $S_NAr$ conditions and subsequent heterocycle formation to afford NAB2 as a white solid (194.1mg, 52% yield). $^1H$ NMR (500 MHz, $CDCl_3$) δ 8.27 (s, 1H), 8.01 (s, 1H), 7.83 (dd, *J* = 8.5, 1.7 Hz, 1H), 7.52 (dd, *J* = 7.0, 2.4 Hz, 1H), 7.40 (dd, *J* = 7.0, 2.2 Hz, 1H), 7.31 (d, *J* = 7.9 Hz, 1H), 7.29 – 7.22 (m, 3H), 7.18 (d, *J* = 8.5 Hz, 1H), 7.11 (s, 1H), 6.71 (bs, 1H), 4.79 (d, *J* = 5.9 Hz, 2H), 2.40 (s, 3H), and 2.03 ppm (s, 3H). $^{13}C$ NMR (126 MHz, $CDCl_3$) δ 167.85, 144.65, 143.01, 137.45, 137.01, 136.00, 134.14, 133.78, 132.04, 131.53, 130.55, 130.37, 129.67, 129.27, 129.03, 128.10, 127.27, 123.49, 119.26, 110.91, 42.24, 20.89, and 17.19 ppm. HRMS (ESI-TOF) m/z calcd for $C_{23}H_{20}ClN_3O$ $[M+H]^+$ 390.1368, found 390.1371.

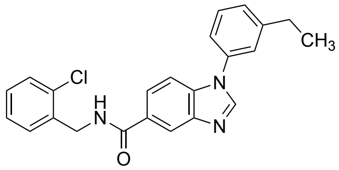

*N-(2-chlorobenzyl)-1-(3-ethylphenyl)-1H-benzo[d]imidazole-5-carboxamide (NAB4).* NAB4 was prepared according to General Procedure A using 2-chlorobenzylamine to generate intermediate *N*-(2-chlorobenzyl)-4-fluoro-3-nitrobenzamide. The benzamide intermediate was purified and carried forward and reacted with 2-ethylaniline under $S_NAr$ conditions and subsequent heterocycle formation to afford NAB4 as a white solid (76% yield). $^1H$ NMR (500 MHz, $CDCl_3$) δ 8.26 (d, *J* = 1.7 Hz, 1H), 8.18 (s, 1H), 7.88 (dd, *J* = 8.6, 1.7 Hz, 1H), 7.57 (d, *J* = 8.6 Hz, 1H), 7.54–7.46 (m, 2H), 7.44 – 7.38 (m, 1H), 7.36 – 7.30 (m, 3H), 7.28 – 7.23 (m, 2H), 6.70 (s, 1H), 4.79 (d, *J* = 6.0 Hz, 2H), 2.78 (q, *J* = 7.6 Hz, 2H), and 1.32 ppm (t, *J* = 7.6 Hz, 3H). $^{13}C$ NMR (126 MHz, $CDCl_3$) δ 166.51, 151.17, 143.91, 138.75, 135.91, 135.77, 135.75, 132.67, 130.51, 130.31, 130.07, 129.63, 129.06, 128.11, 127.22, 123.55, 123.34, 121.38, 119.27, 110.79, 42.25, 28.77, and 15.42 ppm. HRMS (ESI-TOF) m/z calcd for $C_{23}H_{20}ClN_3O$ $[M+H]^+$ 390.1295, found 390.1371.

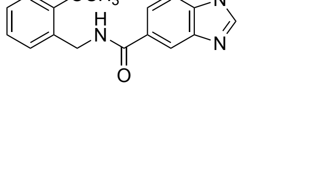

*N-(2-methoxybenzyl)-1-(m-tolyl)-1H-benzo[d]imidazole-5-carboxamide (NAB9).* NAB9 was prepared according to General Procedure B. Briefly, 2-methylaniline (*o*-toluidine) was reacted with 4-fluoro-3-nitrobenzoic acid under $S_NAr$ conditions. The crude nitroaniline was purified and carried forward for amide bond formation with 2-methoxybenzylamine and subsequent heterocycle formation to NAB9 as a white solid (41% yield). $^1H$ NMR (500 MHz, $CDCl_3$) δ 8.20 (d, $J$ = 1.7 Hz, 1H), 8.16 (s, 1H), 7.88 (dd, $J$ = 8.5, 1.7 Hz, 1H), 7.56 (d, $J$ = 8.5 Hz, 1H), 7.50 – 7.43 (m, 1H), 7.39 (dd, $J$ = 7.4, 1.7 Hz, 1H), 7.33 – 7.26 (m, 4H), 6.99 – 6.86 (m, 2H), 6.79 (m, 1H), 4.70 (d, $J$ = 5.8 Hz, 2H), 3.92 (s, 3H), and 2.48 ppm (s, 3H). $^{13}C$ NMR (126 MHz, $CDCl_3$) δ 167.25, 157.76, 143.76, 143.65, 140.47, 135.87, 130.07, 129.95, 129.20, 128.98, 126.35, 124.69, 123.50, 121.15, 120.82, 119.01, 110.67, 110.40, 55.48, 40.30, and 21.44 ppm. HRMS (ESI-TOF) m/z calcd for $C_{23}H_{21}N_3O_2$ $[M+H]^+$ 372.1634, found 372.1709.

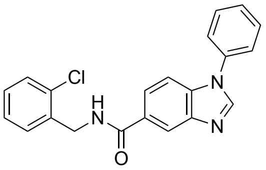

*N-(2-chlorobenzyl)-1-phenyl-1H-benzo[d]imidazole-5-carboxamide (NAB13).* NAB13 was prepared according to General Procedure A using 2-chlorobenzylamine to generate intermediate *N*-(2-chlorobenzyl)-4-fluoro-3-nitrobenzamide. The benzamide intermediate was purified and carried forward and reacted with aniline under $S_NAr$ conditions and subsequent heterocycle formation to afford NAB13 as a white solid (41% yield). $^1H$ NMR (500 MHz, $CDCl_3$) δ 8.27 (d, $J$ = 1.6 Hz, 1H), 8.18 (s, 1H), 7.88 (dd, $J$ = 8.5, 1.7 Hz, 1H), 7.64 – 7.55 (m, 3H), 7.55 – 7.47 (m, 4H), 7.44 – 7.34 (m, 1H), 7.31 – 7.22 (m, 2H), 6.71 (m, 1H), and 4.79 ppm (d, $J$ = 6.0 Hz, 2H). $^{13}C$ NMR (126 MHz, $CDCl_3$) δ 160.67, 143.85, 143.38, 137.53, 135.31, 135.30, 132.19, 130.52, 130.24, 129.63, 129.07, 128.52, 127.22, 124.13, 123.44, 119.32, 110.70, and 42.26 ppm. HRMS (ESI-TOF) m/z calcd for $C_{21}H_{16}ClN_3O$ $[M+H]^+$ 362.0982, found 362.1063.

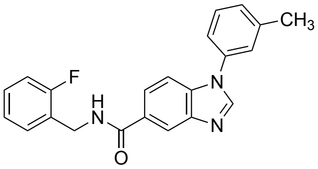

*N-(2-fluorobenzyl)-1-(m-tolyl)-1H-benzo[d]imidazole-5-carboxamide (NAB15)*. NAB15 was prepared according to General Procedure B. Briefly, 2-methylaniline (*o*-toluidine) was reacted with 4-fluoro-3-nitrobenzoic acid under $S_NAr$ conditions. The crude nitroaniline was purified and carried forward for amide bond formation with 2-fluorobenzylamine and subsequent heterocycle formation to NAB15 as a white solid (36% yield). $^1H$ NMR (500 MHz, $CDCl_3$) δ 8.25 (d, $J$ = 1.6 Hz, 1H), 8.16 (s, 1H), 7.87 (dd, $J$ = 8.6, 1.7 Hz, 1H), 7.57 (d, $J$ = 8.5 Hz, 1H), 7.51 – 7.44 (m, 2H), 7.31 (m, 4H), 7.14 (td, $J$ = 7.5, 1.2 Hz, 1H), 7.09 (ddd, $J$ = 9.8, 8.2, 1.2 Hz, 1H), 4.76 (d, $J$ = 5.9 Hz, 2H), and 2.48 ppm (s, 3H). $^{13}C$ NMR (126 MHz, $CDCl_3$) δ 167.56, 161.2 (d, $J$ = 247.1 Hz), 143.91, 142.69, 140.50, 135.99, 135.97, 130.46 (d, $J$ = 4.4 Hz), 129.97, 129.37, 129.32, 129.26,

125.29 (d, $J$ = 5.9), 124.72, 124.44, 124.41 (d, $J$ = 3.6), 123.32, 121.17, 119.28, 115.47 (d, $J$ = 20.9), 110.76, 38.31 (d, $J$ = 4.1 Hz), and 21.44 ppm. HRMS (ESI-TOF) m/z calcd for $C_{22}H_{18}FN_3O$ $[M+H]^+$ 360.1434, found 360.1510.

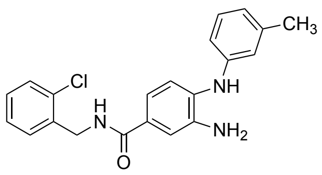

*3-amino-N-(2-chlorobenzyl)-4-(m-tolylamino)benzamide (NAB17).* NAB17 was prepared according to General Procedure A using 2-chlorobenzylamine to generate intermediate *N*-(2-chlorobenzyl)-4-fluoro-3-nitrobenzamide. The benzamide intermediate was purified and carried forward and reacted with 2-methylaniline (*o*-toluidine) under $S_NAr$ conditions. The crude nitroaniline was carried forward into the final step except formic acid was omitted from this reaction mixture to prevent closure of the heterocycle. NAB17 was afforded as an off-white solid (72.7 mg, 62.1%). $^1H$ NMR (500 MHz, $CDCl_3$) δ 7.50 – 7.44 (m, 1H), 7.39 (dd, $J$ = 7.2, 2.2 Hz, 1H), 7.31 (d, $J$ = 2.0 Hz, 1H), 7.25 – 7.20 (m, 2H), 7.17 – 7.08 (m, 3H), 6.74 (d, $J$ = 7.6 Hz, 1H), 6.69 (d, $J$ = 6.2 Hz, 2H), 6.49 (m, 1H), 5.33 (s, 1H), 4.72 (d, $J$ = 6.0 Hz, 2H), 2.29 (s, 3H), and 2.01 ppm (d, $J$ = 1.2 Hz, 1H). $^{13}C$ NMR (126 MHz, $CDCl_3$) δ : 167.30, 143.44, 139.53, 139.46, 135.97, 133.86, 133.81, 130.58, 129.70, 129.56, 129.42, 129.12, 127.32, 121.90, 120.57, 118.06, 117.93, 115.86, 114.44, 42.13, and 21.66 ppm. HRMS (ESI-TOF) m/z calcd for C21H20ClN3O $[M+H]^+$ 366.1368, found 366.1376.

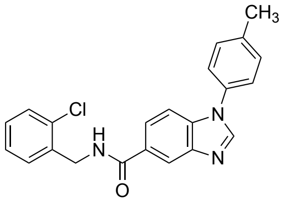

*N-(2-chlorobenzyl)-1-(p-tolyl)-1H-benzo[d]imidazole-5-carboxamide (NAB19).* NAB19 was prepared according to General Procedure A using 2-chlorobenzylamine to generate intermediate *N*-(2-chlorobenzyl)-4-fluoro-3-nitrobenzamide. The benzamine intermediate was purified and carried forward and reacted with 4-methylaniline (*p*-toluidine) under $S_NAr$ conditions and subsequent heterocycle formation to afford NAB19 as a white solid (61% yield). $^1H$ NMR (500 MHz, $CDCl_3$) δ 8.25 (s, 1H), 8.15 (s, 1H), 7.86 (d, $J$ = 8.6 Hz, 1H), 7.53 (m, 2H), 7.43 – 7.37 (m, 5H), 7.32–7.29 (m, 2H), 6.69 (m, 1H), 4.79 (d, $J$ = 5.9 Hz, 2H), and 2.47 ppm (s, 3H). $^{13}C$ NMR (126 MHz, $CDCl_3$) δ 167.43, 143.96, 143.65, 138.67, 137.42, 135.78, 134.19, 133.15, 130.73, 130.50, 129.62, 129.26, 129.05, 127.21, 124.05, 123.31, 119.24, 110.70, 42.25, and 21.17 ppm. HRMS (ESI-TOF) m/z calcd for $C_{22}H_{18}ClN_3O$ $[M+H]^+$ 376.1138, found 376.1215.

### Mammalian tissue culture

SHSY5Y cells were obtained from the Duke University Cell Culture Facility and were maintained at 37 °C with 5% $CO_2$ in DMEM:F12 (1:1; Gibco) supplemented with 10% FBS (Sigma Aldrich) and 1X penicillin-streptomycin (Gibco). Cells were passaged at 80% confluency with a sub-cultivation ratio of 1:20 unless seeding for transfection experiments where a higher density was required. Transfection of SHSY5Y was performed with GeneX Plus (ATCC).

### One-pot TPP/SPROX analysis

*SPROX Analysis.* NAB2 and DMSO treated SHSY5Y lysate samples were subjected to a SPROX analysis similar to that previously described using the one-pot strategy recently reported.[3–6] Five biological replicates of each condition were processed according to the following method. Briefly, aliquots of NAB2 and DMSO treated samples were distributed into a series of 12 PBS buffers (pH 7.4) containing increasing urea concentrations. The five pairs of 12 NAB2 and DMSO samples in the urea-containing buffers generated above were equilibrated at room temperature for 1 h. The methionine oxidation reaction in SPROX was initiated by adding an aliquot of 30% (v/v) $H_2O_2$ to reach the final concentration of ~0.8M, and the oxidation reaction was allowed to proceed for 3 min. The oxidation reaction was quenched by adding a 500 µL aliquot of 500 mM TCEP. For each replicate, equal amounts of the NAB2-treated samples were combined as were equal amounts of the DMSO treated samples, affording 10 total samples. Samples were transferred to centrifugal filters (Millipore Amicon Ultra centrifugal filter units, 10 kDa MWCO) for buffer exchange into 8 M urea in 0.1 M Tris-HCl pH 8.5 followed by 5mM TCEP in 8M urea for 1 h at room temperature, 20mM MMTS in 8M urea for 15 min at room temperature before a buffer exchange to 0.1M TEAB (triethylammonium bicarbonate) and an overnight digestion with trypsin at 37 °C using an enzyme/protein ratio of ~1:100. Digested samples were subjected to TMT10Plex labeling according to the manufacturer's protocol. Labeled peptides were centrifuged through the filters.

*TPP Analysis.* NAB2 and DMSO treated SHSY5Y lysate samples were subjected to a TPP analysis similar to that previously described using the one-pot strategy recently reported.[5–7] Five biological replicates of each condition were processed according to the following method. Aliquots of the NAB2- and DMSO-treated samples were distributed into a series of 12 different tubes prior to thermal denaturation which involved heating the protein material in the NAB2 and DMSO ligand samples for 3 min at one of 12 temperatures that were equally spaced at 2 °C intervals between 43 and 65 °C. The protein samples were removed from the heat and equilibrated at room temperature for 3 min prior to placing them on ice. The NAB2- and DMSO-treated samples were combined to generate a single NAB2 and DMSO sample. The resulting NAB2 and DMSO ligand samples were centrifuged at 48,000 rpm for 20 min using a TPA100.1 rotor and a Beckman Optima TL ultracentrifuge. The supernatant (800 µL) from each combined sample was transferred into a centrifugal filter unit (10 kDa MWCO) for processing for quantitative bottom-up proteomic analysis as previously described[4]. Briefly, sample processing involved TCEP reduction, reaction with MMTS, digestion with trypsin, and labelling with a TMT10Plex reagent kit according to the manufacturer's protocol. Labeled peptides were centrifuged through the filters.

*Multiplexing in preparation for LC-MS/MS analysis.* Equal volumes of solution from each TMT10Plex labeled sample were combined into one tube. A $C_{18}$ Macrospin column cleanup was performed on the combined, labeled sample. For SPROX samples only, samples were subsequently processed by enrichment of methionine-containing peptides with the Pi3 Methionine reagent kit according to manufacturer's protocol. The enriched sample was dried in a speed vac prior to LC-MS/MS analyses.

*LC-MS/MS analysis.* The samples were reconstituted in 2% acetonitrile and 1% trifluoroacetic acid in water to reach a concentration of ~1mg/ml. The sample (1ug aliquot) was injected to the nLC-1200 (Thermo Scientific) and separated on PepMap $C_{18}$ column (Thermo Scientific, 2$\mu$m, 100Å, 75um x 25cm) with a 2 h gradient before analyzed by Thermo Orbitrap Exploris 480 with following parameters: 120000 resolution, 300% AGC target and 2.5s scan cycle for MS1; 45000 resolution, 300% AGC target and 36% HCD collision energy for MS2. The acquired data were searched against *Homo sapiens* proteome (Uniprot ID: UP000005640) acquired from UniprotDB using Proteome Discoverer 2.3 (Thermo Scientific). The searched data were analyzed following the data analysis protocol as described previously.[4] Briefly, five ratios were calculated between ligand/vehicle group, and a two-tailed $t$ test was conducted to calculate the $p$ value used for hit selection.

**Protein thermal shift assay for measurement of ligand-induced changes in protein stability**

Recombinant Rab1a (2 $\mu$M) was pre-incubated with GDP or GppNHp before incubation with NAB2 at various concentrations or NAB derivatives at 50 $\mu$M for 30 minutes. Following incubation, SYPRO orange (5000X stock in DMSO, Invitrogen) was added to a final concentration of 5X and mixtures were aliquoted to a final volume of 15 $\mu$L into a LightCycle 96-well white qPCR plate (Roche) with a minimum of three replicates. Thermal denaturation was conducted in a LightCycler 480 (Roche) via continuous heating from 20 to 85 °C over 18 minutes. Experimental $T_m$ was calculated as the absolute minimum of the negative first derivative curve of the melting curve (i.e. –δ(fluorescence intensity)/δ(temperature)).

**Equilibrium dialysis for measurement of NAB2 binding to Rab1**

Recombinant Rab1 (5 $\mu$M) was prepared in its apo form or in the presence of GDP (10 $\mu$M) or GppNHp (10 $\mu$M) in phosphate buffer (20 mM $Na_2HPO_4$, 50 mM NaCl, 2 mM $MgCl_2$, pH 7.4). Samples were treated with NAB2 (15 $\mu$M) or vehicle control with a final DMSO concentration of 1.5% to provide a final solution of 50 $\mu$L in total volume. Additionally, positive controls (NAB2 only without Rab1) and a negative control (Rab1-apo only, no NAB2) were assembled. All samples were prepared in triplicate and loaded into one chamber of DispoEquilibrium dialysis cassettes (10 kDa MWCO, Harvard Apparatus). The other chamber of each cassette was loaded with 50 $\mu$L of buffer control (20 mM $Na_2HPO_4$, 50 mM NaCl, 2 mM $MgCl_2$, pH 7.4 with 1.5% DMSO). Samples were kept level and incubated with gentle orbital agitation at 4 °C for 24 hours. Following equilibration, the buffer chamber sample of each cassette was collected and samples were analyzed by RP-HPLC-MS via Agilent 6460 Triple Quadrupole LC-MS (a 2.6 µm EVO–$C_{18}$,

100×3 mm column) using a linear gradient of 25–75% B over 8 minutes, where mobile phase A was 100: 3:0.3 $H_2O$/MeOH/TFA and mobile phase B was 100: 3:0.3 MeCN/$H_2O$/TFA. ESI-MS was performed in positive ion mode and total ion chromatograms (TICs) were extracted using a m/z charge filter of 390 ± 0.5, to yield the extracted ion chromatogram (EIC) corresponding to the $[M+H]^+$ adduct of NAB2. AUC of each EIC was determined by integration of the EIC peak with a retention time of 4.8 to 5 minutes. AUC of all samples were analyzed using Prism GraphPad and statistical analyses were performed to compare all samples to the positive control using an unpaired t-test with Welch's correction for unequal variances.

**Malachite green activity assay for Rab1-dependent GTP consumption**

Rab1a activity was monitored using a malachite green assay according to the manufacturer's recommended protocol. The assay was obtained from Sigma-Aldrich (catalog no. MAK307). Recombinant Rab1 (5 $\mu$M) was incubated with GTP in the presence of NAB2 (0 – 100 $\mu$M) with GTP (1 mM) in Rab1a assay buffer (20 mM Tris, 150 mM NaCl, pH 7.5) for up to three hours before quenching with malachite green developing reagent. Quenched reactions were incubated for 30 minutes and $A_{620}$ was determined using the SpectraMax iX3 plate reader. All samples were background subtracted, and phosphate concentration was determined by comparison to a phosphate standard curve as described in the recommended protocol. Percent activity was calculated as phosphate consumption in the presence of NAB2 treatment relative to a DMSO control (containing no NAB2). Data was visualized with Prism GraphPad.

**Subcellular fractionation of SHSY5Y cells**

SHSY5Y cells were seeded into 6-well plates. At 24h post seeding, cells were treated with DMSO control or NAB2 (20 $\mu$M) for 12 hours. Following treatment, proteins were harvested from the cells by subcellular fractionation using ProteoExtract subcellular fractionation kit (Millipore Sigma). Fractionated samples were analyzed by SDS-PAGE and electrophoresis for detection of Rab1a (anti-Rab1a, 1:2000, ThermoFisher no. 11671-1-AP) and actin (anti-actin, 1:2000, Abcam ab12148). Rab1a signal was quantified by ImageJ,[8] normalized to actin loading control signal, and plotted as percentage of total normalized Rab1a signal. Data was processed using Prism GraphPad.

**Confocal immunofluorescence microscopy**

SHSY5Y cells were seeded for analysis in 6-well plates with pre-treated cover slips. Twenty-four hours post seeding, cells were treated with NAB2 (20 $\mu$M) or DMSO control for 12h. Following treatment, cells were fixed using cold methanol and blocked with 3% BSA in TBST for 30 minutes. Following blocking, cells were treated with anti-Rab1a (ThermoFisher no. 11671-1-AP) primary antibody, Rab GDI primary antibody (Santa Cruz Biotechnology, sc-374649), and Concanavalin A (AlexaFluor 594 conjugated, Fisher Scientific C11253) for one hour and then with species-specific AlexaFluor conjugated secondary antibodies and Hoechst stain. Cover slips were fixed on microscopy slides and analyzed using a Zeiss AiryScan 880 confocal microscope using the 40X oil objective. Images were processed using Imaris software. Quantitation of co-localization was determined by defining single as regions of interest (ROI) and calculating the

Pearson's Correlation Coefficient (PCC) across the full volume (z-stack) of the ROI. A minimum of 15 ROI across three separate microscopy frames were measured for each condition and PCC was reported as average ± s.e.m. for comparison of NAB2 treated (6 hr post-treatment) vs time = 0 control.

**MTT cell viability assay**

Twenty-four hours pre-transfection, SHSY5Y cells were seeded into 6- or 24-well tissue cultures plates using the described culturing conditions. Three hours pre-transfection, cells were treated with fresh media. Cells were subsequently co-transfected with equal amounts of pHM6-α-synuclein-A53T (Addgene #40825) and pCMV-Rab1a (Addgene #46776), pCI-HA-Nedd4 (Addgene #27002) or empty vector control using GeneX Plus transfection reagent according to manufacturer's protocol. Twenty-four hours post-transfection, media was refreshed. At forty-eight hours post-transfection, cells processed by MTT viability assay or treated with NAB2 as described prior to MTT analysis.

The MTT assay protocol was adapted from the recommended protocol generated by Abcam, where culture media is removed from cells and cells were treated with serum free media and MTT (5 mg/mL in PBS) in a 1:1 ratio and incubated for three hours. The assay was then quenched and insoluble formazan crystals generated by cellular reduction of MTT during the incubation time is subsequently dissolved with the addition of 4mM HCl in isopropanol with 0.1% NP-40 detergent. Quenched plates were incubated at rt for 15 minutes and absorbance was measured in a SpectraMax iX3 plate reader using an endpoint assay protocol with absorbance wavelength set to 590 nm. Absorbance of replicate samples was used to calculate average absorbance for each condition and data was processed with Prism GraphPad.

### Supplemental Figures

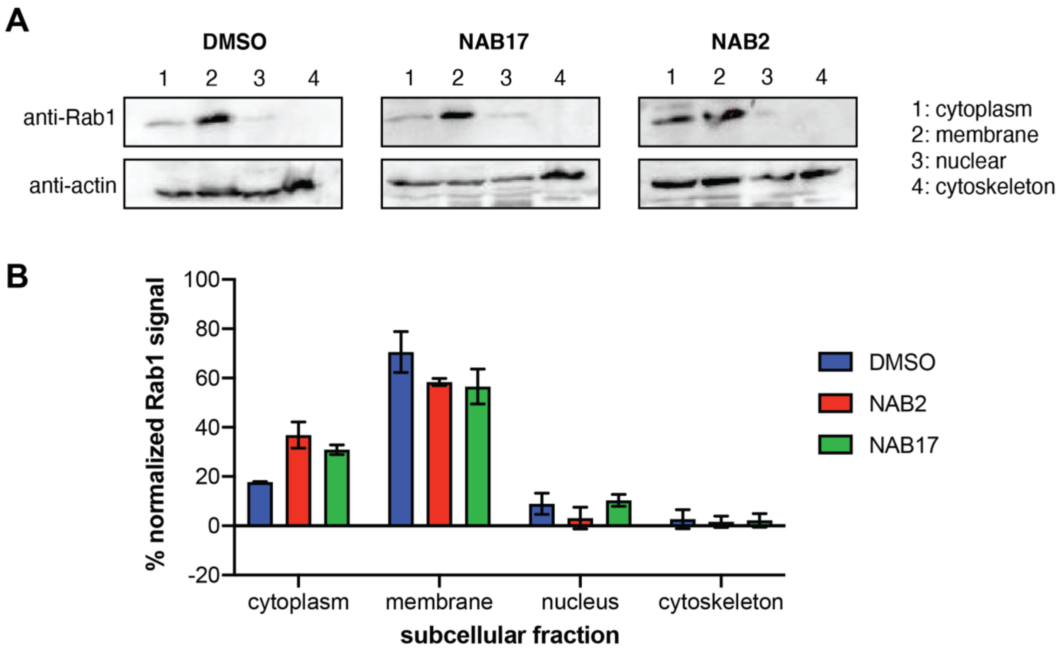

**Figure S1: Rab1 localization was monitored in a NAB2-dependent manner via subcellular fractionation.** Rab1a compartmentalization in SHSY5Y cells was **(A)** monitored through subcellular fractionation (ProteoExtract, Millipore Sigma) and detected via immunoblotting. Compartmentalization was monitored in a NAB2- and NAB17-dependent manner, and **(B)** chemiluminescent signal intensity was quantified and normalized to anti-actin signal as a loading control. Quantification reveals no significant NAB2-dependent alteration of Rab1 compartmentalization. Samples were prepared and analyzed in triplicate, with representative blots shown in **(A)** and data in **(B)** presented as mean ± s.e.m. of triplicate experiments. Immunoblots developed using chemiluminescent ECL reagents and images processed using ImageJ.[8]
